# Polygenic risk score prediction accuracy convergence

**DOI:** 10.1101/2023.06.27.546518

**Authors:** Léo Henches, Jihye Kim, Zhiyu Yang, Simone Rubinacci, Gabriel Pires, Clara Albiñana, Christophe Boetto, Hanna Julienne, Arthur Frouin, Antoine Auvergne, Yuka Suzuki, Sarah Djebali, Olivier Delaneau, Andrea Ganna, Bjarni Vilhjálmsson, Florian Privé, Hugues Aschard

## Abstract

Polygenic risk scores (PRS) trained from genome-wide association study (GWAS) results are set to play a pivotal role in biomedical research addressing multifactorial human diseases. The prospect of using these risk scores in clinical care and public health is generating both enthusiasm and controversy, with varying opinions about strengths and limitations across experts^1^. The performances of existing polygenic scores are still limited, and although it is expected to improve with increasing sample size of GWAS and the development of new powerful methods, it remains unclear how much prediction can be ultimately achieved. Here, we conducted a retrospective analysis to assess the progress in PRS prediction accuracy since the publication of the first large-scale GWASs using six common human diseases with sufficient GWAS data. We show that while PRS accuracy has grown rapidly for years, the improvement pace from recent GWAS has decreased substantially, suggesting that further increasing GWAS sample size may translate into very modest risk discrimination improvement. We next investigated the factors influencing the maximum achievable prediction using recently released whole genome-sequencing data from 125K UK Biobank participants, and state-of-the-art modeling of polygenic outcomes. Our analyses point toward increasing the variant coverage of PRS, using either more imputed variants or sequencing data, as a key component for future improvement in prediction accuracy.

## Main

Most common human diseases harbor a strong polygenic inheritance, characterized by a very large number of genetic variants of small effects. This scattered distribution of risk has severely hampered the initial goal of using genetic association studies for personalized medicine through individualized disease risk predictions, prevention strategies and treatments ^2-4^. This issue has been recognized early on in the GWAS era, and the community has developed a strong case for genetic risk profiling on the basis of polygenic risk scores (PRSs), derived from genome-wide association studies (GWAS) results^5^. In its simplest form, a PRS for an individual is a summation of multiple single nucleotide polymorphisms (SNPs) weighted by their effect sizes estimated from independent GWAS data. Initially PRSs were constructed from a small number of independent genome-wide significant variants, but they have evolved to include thousands to millions of variants selected from the complete GWAS results, using optimized selection criteria and weighting schemes^6,7^. As for predicting any highly multifactorial outcome, the accuracy of PRS predictions mostly depends on the sample size of the dataset used to estimate individual variant effects. However, despite a substantial increase in GWAS sample size over time, the predictive performance of existing PRSs remains low, raising concerns about their value for clinical purposes^8-10^.

Several studies have estimated the potential for genetic risk prediction in polygenic diseases describing links between prediction accuracy, typically defined by the area under the receiver operating characteristic curve (AUC), disease prevalence and heritability^11-13^. More recent studies also proposed theoretical frameworks to estimate future improvement in PRS-based prediction based on the distribution of effects at causal variants^14,15^. However, none of those examined performances retrospectively. This is partly explained by a lack of standard in the implementation of PRS, and the impact of various factors inducing heterogeneity in the PRS performances^16,17^. Here we examine how PRS prediction accuracy of six diseases evolved during the past 15 years and how existing models, state-of-the-art sequencing data, and functional annotation data can inform potential future improvements. We use only PRS derived from population of European ancestry. Using PRS derived in non-European population would be challenging because of the current lack of large-scale GWAS data. However, the linear relationship observed in cross-ancestry PRS transferability^18,19^ suggests the trends should remain consistent across populations.

### PRS prediction accuracy and sample size

We first pulled from the literature and curated previous reports of genetic risk score prediction accuracy, expressed as the AUC, for coronary artery disease, breast cancer, type 2 diabetes, Alzheimer disease, asthma, and obesity (**Supplementary Note** and **Table S1**). The studies spanned from 2006 to 2020 and the effective sample size *N*_*eff*_ ranged from 981 to 339,224. The reported predictive power showed a very modest association trend with sample size and was instead characterized by substantial heterogeneity (**Figure 1a**). AUC increase was nominally significant for breast cancer (*β* = 3.3 ×10^−4^ per 1K increase in sample size, *P*=0.018) and almost nominally significance for type 2 diabetes (*β* = 3.9×10^−4^, *P*=0.091) and obesity (*β* = 2.2×10^−4^, *P*=0.078), but not significant for the other outcomes, with for example, a negative trend for coronary artery disease (*β* = -6.8×10^−5^, *P*=0.73) despite an effective sample size varying from *N* _*eff*_ =4,522 to *N*_*eff*_ =184,305. This is likely due to the limited number of studies available, but also to a number of factors already discussed in the literature and complex to decipher, including heterogeneity in the characteristics of the population (age, sex, fine-scale genetic ancestry within European population), disease definition, and the method used to derive the PRS weights^16,20^.

**Figure 1.**
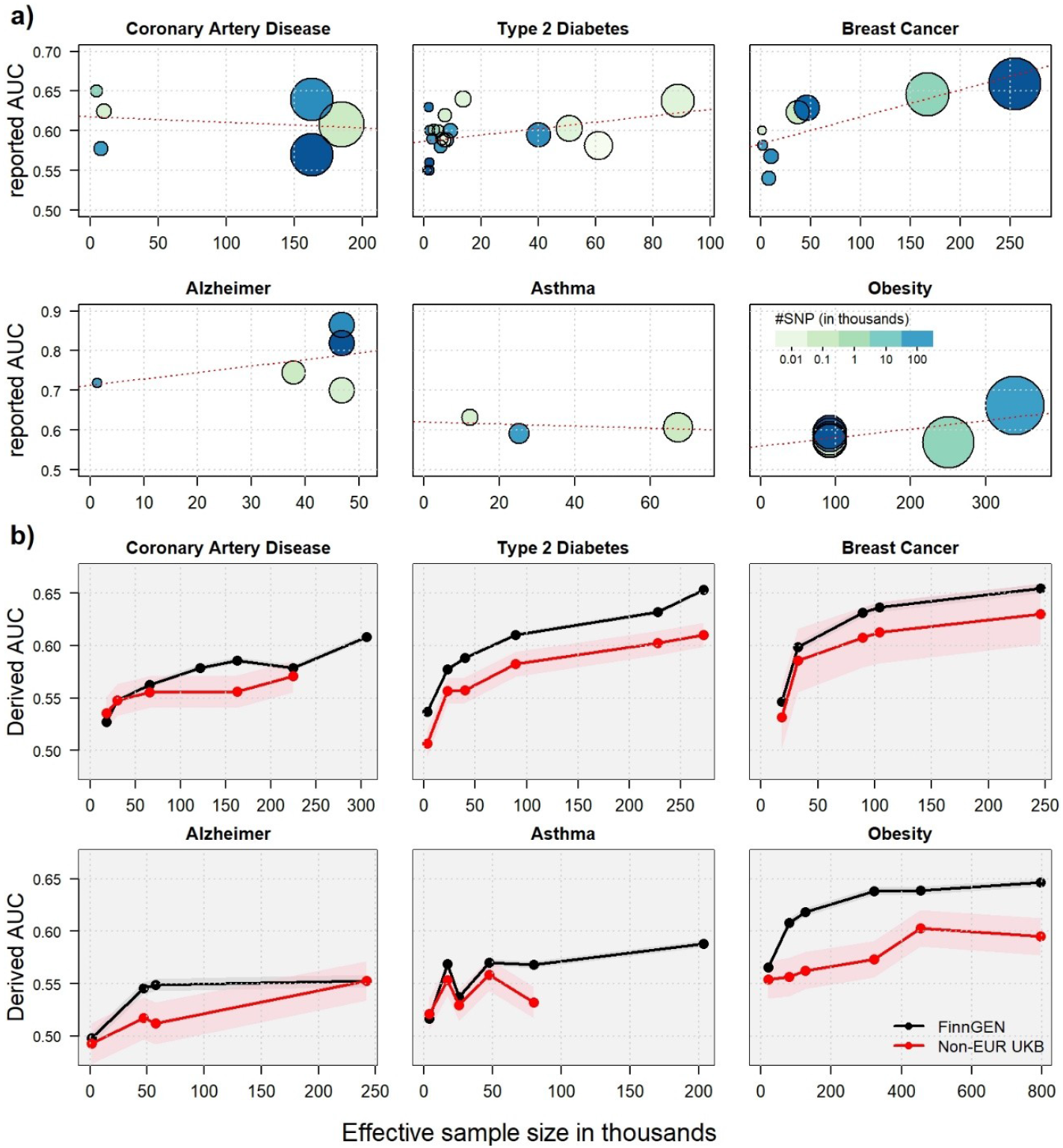
Polygenic risk scores predictive accuracy as a function of sample size. Panel a) presents the area under the receiver operating characteristic curve (AUC) reported in the literature for polygenic risk score as a function of the effective sample size for coronary artery disease, type 2 diabetes, breast cancer, Alzheimer disease, asthma and obesity. The color gradient corresponds to the number of variants used, from a few top associated variants (light green) to millions (dark blue), and the size is proportional to the log of the effective sample size. Panel b) shows AUCs for the six outcomes derived using a harmonized pipeline. Polygenic risk scores were trained from 34 genome-wide association study (GWAS) using the LDpred2 approach, and tested in individual-level data from the FinnGEN cohort (black) and in six non-European ancestry *UK Biobank* populations (red, meta-analysis over six populations). The AUCs are plotted as a function of the effective sample size of the corresponding GWAS. Missing values correspond to cases where there was a sample overlap between the test and train sets.

To quantify more precisely changes in the predictive accuracy as a function of sample size, we implemented a harmonized pipeline to derive PRS and applied it to a set of highly curated GWAS data. We derived PRS weights for the six outcomes using 29 GWAS summary statistics (with different sample sizes) published since 2007 and including mostly participants of European ancestry. To expend the sample size coverage, we also added five GWAS with modest sample size that we conducted using unrelated participants of European ancestry from the UK Biobank cohorts^21^ (**Supplementary Note** and **Table S2**). Those 34 PRS were used to derive AUC in two independent cohorts. Because almost all existing genetic cohorts of European ancestry have been used to produce these GWAS, we focused our application on relative power and investigated the trend in AUC in 16,000 participants from six different non-European ancestries from the *UK Biobank* cohort, and up to 392,423 participants of Finnish ancestry from the FinnGen cohort^22^ (**Table S3**). Difference in genetic ancestry induces lower absolute predictive performance, however, because the PRS portability across population is expected to be linear^18,19^, the trend remains highly informative, and for example, top GWAS hits between FinnGen and European ancestry samples were highly concordant (average squared-correlation between effect estimates equals 0.56, **Fig. S1**).

The prediction accuracy shows a univocal non-linear increasing trend as a function of sample size that starts with a sharp increase for the few first studies followed by a gradual decline in improvement. The convergence patterns are highly concordant between FinnGen and non-European UK Biobank participants, only displaying the expected offset due to genetic ancestry difference (**Fig 1b, Fig S2-S3**, and **Table S4**). It suggests that PRSs based on future GWASs of the target outcome with larger sample size are unlikely to dramatically improve prediction accuracy (as measured by the AUC) by themselves. The AUC convergence is especially striking for type 2 diabetes, obesity, breast cancer and coronary artery disease. Asthma displays a noisier trend, potentially reflecting the challenges in the disease definition and diagnosis. Alzheimer disease (AD) shows a continuous increase in AUC for the non-European UKB samples, but no increase for the last and largest GWAS in FinnGEN despite a five-fold increase in the effective sample size. This might be explained by the use of a proxy for disease status (a score determined by the Alzheimer status of the parents’ participants) for the derivation of the PRS weights in the largest GWAS study. AD-by-proxy was also used in the non-European UKB participants because of a limited number of cases, while the true AD status was used in FinnGEN (**Supplementary Note**). As recently discussed, cohorts with heterogeneous disease definition and study design can induce variability in heritability estimates of AD^23^. This likely induces reduced prediction in FinnGEN in our analysis.

### Modelling AUC convergence

The convergence of the AUC toward its maximum can be investigated using simulation, and one can easily demonstrate the validity of simple theoretical models^13^ (**Fig. S4**). Predicting the convergence rate in real data is much more challenging and requires the estimation of multiple parameters of the disease genetic model. It includes the heritability, the polygenicity, the distribution (or mixture of distributions) of causal genetic effects, and the dependence of those effects on the linkage disequilibrium, functional annotations, and the minor allele frequency (MAF). We first examined to what extent existing tools are able to fit the AUC derived in real data using the GENESIS approach^14^. Briefly, GENESIS infers some, but not all, of the aforementioned parameters and uses them to predict the expected AUCs as a function of future increases in sample size. As recommended by the authors, we focused on a three-component mixture of effects, where variants are classified as either non-causal, causal with low effect, or causal with large effect. The shape of the predicted AUC as a function of sample size obtained from GENESIS is reasonably close to the ones observed in FinnGen and the UK Biobank (**Fig S5a**), confirming the observation of a sharp decline in PRS predictive power improvement. However, the increase rate and expected maximum AUC varied substantially conditional on the input GWAS used, being especially variable for Alzheimer disease, asthma, and obesity, and highlighting the importance of conducting real data retrospective analysis.

We next investigated to what extent the observed convergence in AUC might be explained by heterogeneity arising when meta-analyzing an increasing number of studies. Heterogeneity in genetic ancestry, disease definition, and environment across the cohorts meta-analyzed, might reduce heritability and increase polygenicity, and ultimately lessen the benefits of larger sample size GWAS for predictive purposes. To assess these hypotheses, we examined the variability in AUC derived using the same procedure as in **Figure 1b**, and disease parameters estimated using SBayesS^24^, while increasing GWAS sample size, but using the same population for the GWAS and parameter estimation –thus providing a benchmark of the behavior when potential heterogeneity is minimized. We used participants of European ancestry from the UK Biobank (referred further as the EUR-UKB experiment), focusing on obesity and using GWAS of body mass index, one of the sole scenarios allowing to achieve reasonably large sample size within a single homogeneous cohort. While the role of genetic heterogeneity cannot be ruled out, we did not find any strong evidence in favor of an effect of heterogeneity. The polygenicity estimate tends to decrease first (potentially due to limited sample size) until it reaches a minimum and then starts to increase for both the EUR-UKB (**Fig. S6a**) and most of the six outcomes GWASs (**Fig. S7a**). Except for the few estimates derived using a limited sample size, heritability tends to be fairly consistent, although much less variable in EUR-UKB (**Fig. S6b**) than across the GWASs (**Fig. S7b**). Estimates of alpha, the minor allele frequency (MAF)-effect size parameter, also display a similar pattern across analyses (**Fig. S6c, Fig. S7c**). Overall, the AUC from the EUR-UKB analysis appears to follow the same trend as the ones derived in FinnGen and non-European UKB participants (**Fig. S6d**).

As a general observation, the disease parameters estimated from both GENESIS^14^ and SBayesS^24^ vary substantially conditional on the input GWAS used (**Figs. S5b-c, S6**), with confidence interval often not overlapping across GWASs from the same outcome. For comparison purposes, we applied three additional methods, sumHer^25^, LDSC^26^, and MiXeR^27^, to all available GWAS with an effective sample size larger than 5,000 (**Table S5**). With a few exceptions, cross-method heritability estimates from the same GWAS tend to be less variable than within-method estimates from GWASs of the same outcome. Polygenicity varies extensively across methods, each of them displaying a different trend as a function of sample size. The reasons for this variability are unclear, and until progresses are made in estimating those and other parameters underlying the distribution of genetic effect, our ability to develop an accurate modelling of the convergence rate of PRS predictive power will likely remain limited, again, highlighting the importance of conducting real data retrospective analyses.

### Maximal achievable prediction

The maximum achievable AUC relies on less parameters than the convergence rate. A central feature to derive this maximum is the variant coverage, which itself determines the proportion of heritability captured by the genetic variants used in the PRS. Existing polygenic risk prediction models rely heavily on genome-wide genotyping studies, thanks to a high cost-genomic coverage ratio^28^ allowing to generate genotype data for very large cohorts, as compared to sequencing studies. This implies that even reaching an infinite sample size, the total genetic variance captured by GWAS, and therefore the maximum prediction accuracy, is bounded by the sub-sampling of all existing variants. How much of the narrow-sense heritability (i.e. the linear additive genetic variance^29^) can be captured by GWAS and how much more will be captured using sequencing data has been previously explored using both theoretical model and whole-exome sequencing data of modest size as compared to GWAS studies^30-35^. Although the size of existing genome sequencing data remains limited and does not allow to fully address this question, some recent sequencing efforts provide opportunities to revisit this question. Here, we used individual-level whole-genome sequencing data from 503,195 variants on chromosome 22 measured in 125,152 participants of European ancestry from the UK Biobank study, to quantify the proportion of heritability captured by genotyped variants under various alternative genetic models.

We assumed the relationship between effect size and variants characteristics follow the so-called alpha model –where the expected effect of a variant is proportional to its variance power *α* ^33,36^ (see **Methods**)– and derived two metrics: i) 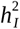, the additive genetic variance captured by both genotyped variants and variants imputed using advanced methods based on haplotype inference^37^, and ii) 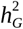, the additive genetic variance captured by genotyped variants and the additional variance captured through linkage disequilibrium between genotyped and untyped variants. In theory, the later parameter should be approximately similar to the GWAS-based heritability 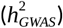 estimated by existing software^38^. Both metrics were derived using sequence data, after considering various alternative approaches (**Fig. S8-S9**). As shown in **Figure 2a**, assuming a random distribution of genetic effect in the genome, 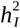 varies from 29% for *α* =− 1.5 to 96% for *α* =0. The heritability captured by genotyped variants only is substantially smaller, with 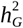 varying from 5% for *α* =− 1.5 to 90% for *α* =0. This suggests that imputed variants with modest and poor imputation quality, often filtered out in GWAS studies, might capture a substantial share of the total narrow-sense heritability missed by genotyped variants. Of note, previous studies argued that the alpha model might overestimate the effect of rare variants^36^. To address this possible limitation, we also devised an attenuated alpha model implying a reduced contribution of rare variants (**Supplementary Note, Fig. S10**). However, when comparing those attenuated models against the standard in real data, we did not find any evidence for an increased fit (**Fig. S11-12**).

**Figure 2.**
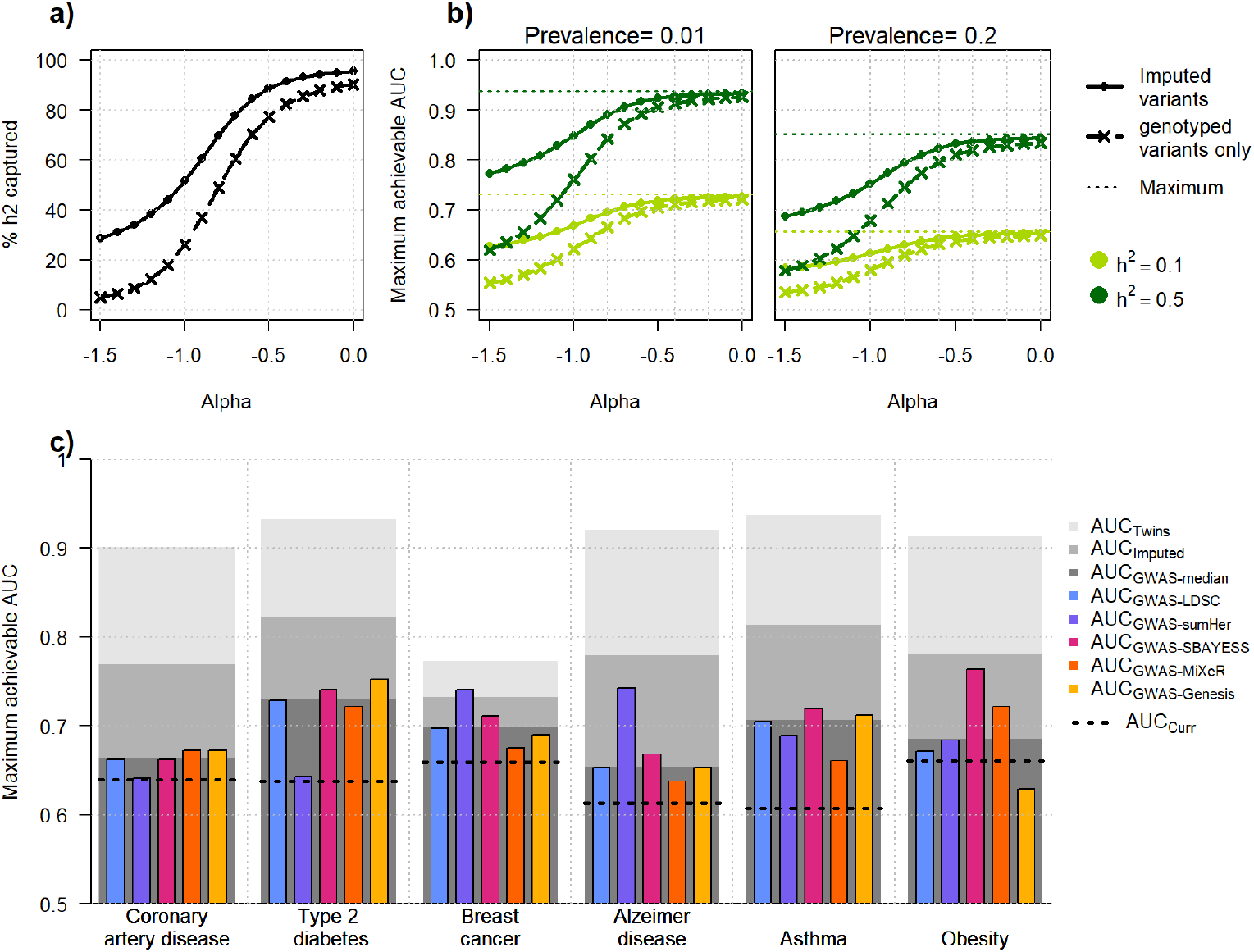
Heritability captured and maximum achievable AUC. Panel a) shows the expected proportion of heritability captured by genotyped (dashed line) and all imputed variants (plain line). Individual-level data from the UK Biobank were use to estimates the squared-correlation between sequenced variants and either genotyped (*ρ*^2^) or imputed variants (*r* ²), and genetic effects were assumed to be distributed following an alpha model. Panels b) shows the corresponding maximum achievable AUC for heritability of 0.1 and 0.5, and disease prevalence equals to 0.01 and 0.2. Panel c) shows estimates of the maximum achievable AUC for six outcomes: coronary artery disease, type 2 diabetes, breast cancer, Alzheimer disease, asthma, and obesity (using body mass index GWAS). Those AUCs were derived based on US disease prevalence pooled, and various estimates of heritability: twin studies (AUC_Twin_), estimates of GWAS heritability derived using five competitive approaches: LDSC regression, sumHer, SBayesS, Genesis, and MiXer (AUC_GWAS_), and twin study heritability captured by imputed variants (AUC_Imputed_). The black dashed lines correspond to the most recent estimates of AUC in real data from the literature, derived based on approximately 1M hapmap3 variants. For Alzheimer disease, heritability estimates and current AUC estimates correspond to a model excluding the APOE region.

Estimates of heritability captured can be translated into maximum achievable AUC given the prevalence and heritability of the target outcome (see **Methods**). As expected, this maximum increases with the disease heritability captured, and decreases with lower value of alpha (**Fig. 2b**). We compared maximum AUC for each of the six diseases using three heritability estimates: twin studies heritability (AUC_twin_), GWAS-based heritability derived using five approaches (AUC_GWAS_, **Table S5-S6**), and heritability captured by all imputed variants (AUC_Imputed_). Note that the latter AUC requires an alpha value. We attempted to use estimates of alpha obtained from various approaches (**Table S6**), but none of them match the expected based on GWAS-based AUC, and we ultimately set alpha for each disease so that the AUC from the genotype-based alpha model equals the median of AUC_GWAS_ across the five approaches (see **Methods**). The comparison of AUC is summarized in **Figure 2d**. First, the AUC_GWAS_ are relatively close to the AUCs reported in recent studies (AUC_curr_, **Table S6**) for coronary artery disease, breast cancer, Alzheimer disease and obesity, confirming that increasing sample size of GWAS for those outcomes is unlikely to dramatically impact prediction performances. Conversely AUC_curr_ for T2D and asthma display sizeable gaps with AUC_GWAS_, so that, despite the convergence observed in **Figure 1b**, future GWAS with larger sample size might provide a slow but continuous improvement in prediction. Second, the gap between AUC_Twin_ and AUC_GWAS_ is large for all outcomes, except for breast cancer, suggesting that increasing the variant coverage in future studies can dramatically improve the predictive power of polygenic risk scores. Third, AUC_Imputed_ is substantially higher than AUC_GWAS_ for all outcomes except breast cancer. Hence, future PRSs using an increasing number of imputed variants, even those with poor imputation quality, have the potential to boost prediction power without requiring the generation of costly sequencing data.

### Effect distributions and predicted performances

In the previous analyses, we assumed that causal variants are randomly distributed across the genome and only modelled the relationship between the effect size and the minor allele frequency. There is strong evidence that causal variants are highly enriched in some functionally annotated regions^39-42^. However, the exact link between those functional annotations and causal variants is not fully understood and likely depends on the outcome study. Rather than assuming a specific relationship, which will be subjective given the current knowledge, we assessed the impact of unequal distribution of causal variants in the genome by estimating the association between a range of functional annotations and the quality of imputation (*r* ², the squared-correlation between sequenced and imputed variants used in the derivation of 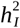). Here we used a total of 1,099 annotations from nine sources (**Table S7-8, Fig. S13**), including gene elements (coding DNA sequence, exon, intron, transcription start and termination sites), DNase I hypersensitive sites (DHS), enhancers and promoters. Those annotations cover 0.001% to 64.7% of the whole genome (**Table S8**).

The largest associations between annotation and *r* ² were observed for gene elements (**Fig. S14a**). Coding DNA sequence (CDS), that cover 1.9% of chromosome 22, show a highly significant decrease in *r* ² (*P* = 1.8 × 10^−307^), with an average *r* ² of 0.27 and 0.40 for CDS and non-CDS variants, respectively. Exons, that cover 9.8% of the chromosome 22, are also strongly negatively associated (*P* = 2.0 × 10^−131^) with an average *r* ² of 0.37 and 0.40 for exonic and non-exonic variants (**Fig. 3a**). Variants from other annotation categories show significant negative and positive associations, but to a lower extent. Enhancers, super enhancers, DHS, and transcription factor binding sites tend to display lower *r* ² than the average of the genome (**Fig. 3b**). Conversely, genetic variants within promoters appear to be slightly better captured. Excluding gene elements, the strongest association was observed for transcription start site with an average *r* ² of 0.36 and 0.40 for annotated and non-annotated variants (*P* = 2.1 × 10^−14^). We did not observe any enrichment for specific cell types or tissues among other top significant annotations (**Fig. S15**).

**Figure 3.**
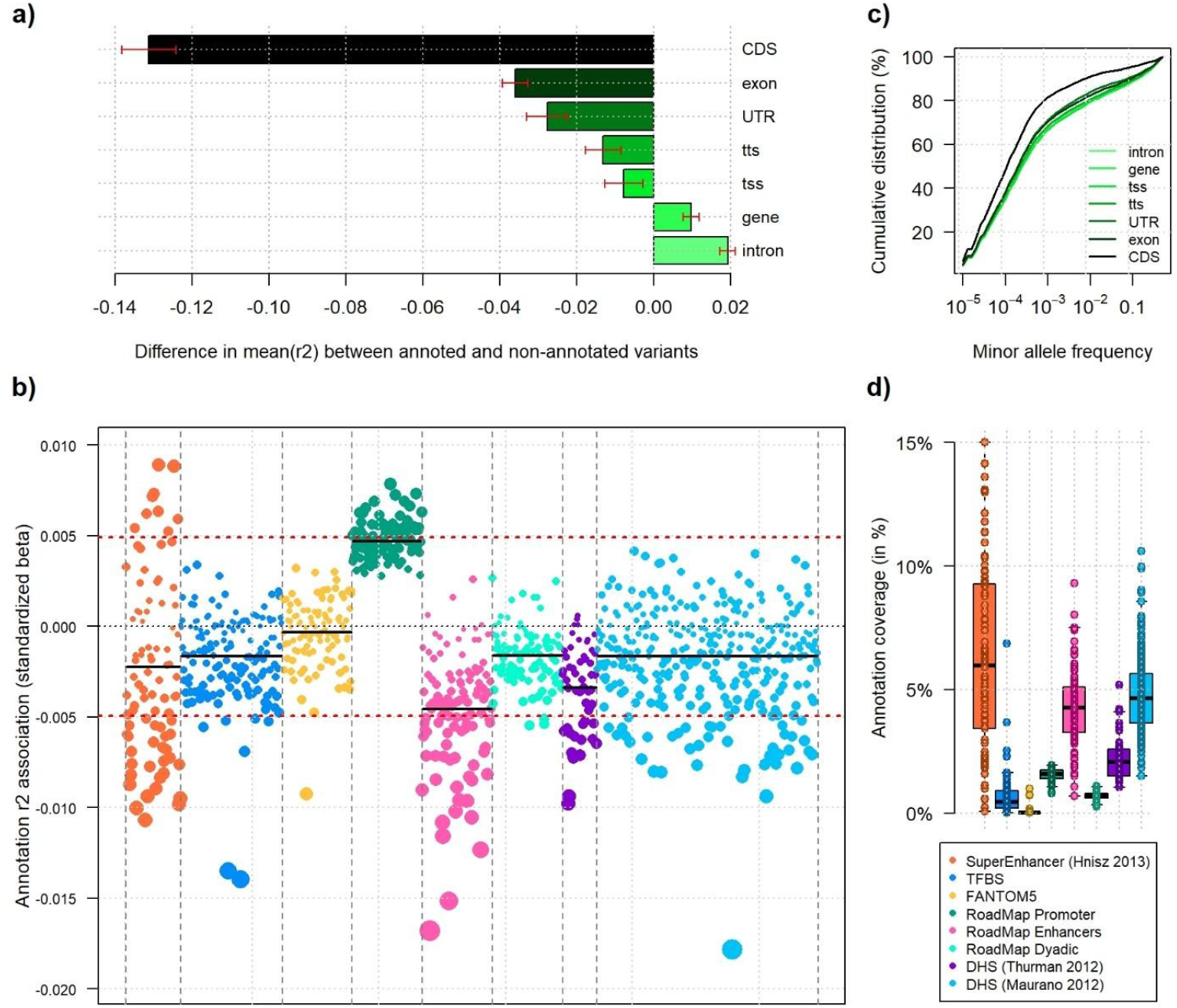
Variability in imputation quality across functional annotations. Relationship between functional annotations and the quality of imputation measured as the squared-correlation (*r* ²) between true and imputed genotypes. Panel a) displays the change in the average *r* ² of chromosome 22 for GENCODE annotations: intron, gene, exon, CDS (coding DNA sequence), tss (transcription start site), tts (transcription termination site), and UTR (untranslated regions). Red bars indicate ±2 standard deviation from the mean and include 95% of the annotated regions. Panel b) displays standardized regression coefficients from univariate association between the measured *r* ² and each of the 1,092 functional annotations from eight categories: TFBS (transcription factor binding site), FANTOM5 (functional annotation of the mammalian genome version 5) regulatory regions, promoters, enhancers, and dyadic from Roadmap, DHS (DNase I hypersensitive sites) derived from two studies, and super enhancers. Horizontal black lines indicate the average per category, and the red dashed lines indicate the significance threshold after correction for multiple testing. Panel c) shows the cumulative distribution of variants for each GENCODE category as a function of the minor allele frequency. Panel d) shows the distribution of frequencies of each annotation, grouped by category.

Part of the decrease in variant coverage (**Fig. 3a-b**) is likely explained by an enrichment for rare variants with, for example, coding regions displaying a significantly higher proportion of rare variants (*P* = 2.5 × 10^−270^, **Fig. 3c** and **Table S9**). However, other annotations positively associated with *r* ² also display a significant enrichment for rare variants (e.g. intron, *P* = 1.0 × 10^−8^), so that minor allele frequency alone is unlikely to fully explain the observed lower variant coverage. As discussed in previous works, causal variants might display decreased linkage disequilibrium with other variants because of selective pressure^34,41^, an assumption that is now commonly incorporated in disease parameter estimation tools^43^. Lower linkage disequilibrium implies lower *r* ², and the observed lower variant coverage for CDS, enhancers, DHS and transcription factor binding sites might embody an enrichment for rare causal variants for those annotations. Under such scenario, the estimated maximum achievable AUCs from imputed variants presented in **Figure 2** could be slightly overestimated, which suggest again that future improvement in PRS prediction accuracy is likely going to rely on improving the variant coverage of future genetic association study (**Fig. S14b**).

## Discussion

Polygenic risk prediction has the potential to transform the diagnosis, treatment, and prevention of many common human diseases. However, the timescale and extent of this transformation is partly unknown, and some in the community continue to express concerns about the relevance of PRS. These concerns are partly due to inconsistencies in the reported performances of polygenic risk models. Previous reports of PRS prediction accuracy are not always properly documented, and while guidelines have been proposed^16^, adherence to reporting statements remains limited. Heterogeneity in the methods used, the populations studied, and the covariates accounted for, can prevent the replication of results and a formal assessment of PRS performances. As illustrated in this study for six outcomes, even a careful curation of publication results based on the above criteria might not be sufficient, and the potential relationship between performances and study parameters can remain blurry.

Here, we demonstrate that the relationship between prediction accuracy of polygenic models and sample size is unequivocal and highly replicable when using harmonized data pre-processing and analysis pipelines. These analyses also show that the prediction accuracy of PRS derived from existing GWAS have started to converge for most diseases studied. As a result, expanding the current state-of-the-art –i.e. fully relying on an increase in GWAS sample size– might only lead to a modest increase in predictive power. For some outcomes, such as coronary artery disease, breast cancer, and obesity, our analyses suggest that the predictive power of GWAS-based PRS is nearing the expected maximum derived based on the GWAS heritability. If confirmed, this implies that improvements in methods used to derive PRS will not either dramatically impact prediction accuracy. For type 2 diabetes and asthma, the gap with the expected maximum remains fairly large, and AUC from future larger GWAS might continue to increase, although any gain will likely require extremely large sample size. Importantly, the observed convergence might be partly explained by increasing heterogeneity arising from large meta-analysis, with for example, very large meta-analysis using a loosen disease definition or increasingly genetically diverse populations, in order to allow for a broader inclusion of participants. Although we did not find any strong evidence for such effect across outcomes we studied, the potential impact of genetic ancestry^18,19^ and phenotype definition^23,44^ heterogeneity is well established. Note that there are other sources of heterogeneity that we did not investigate including in particular sample ascertainment, which might influence the disease prevalence and heritability, and therefore the prediction accuracy.

Estimating the trend in prediction accuracy as a function of sample size depends on our ability to estimate for each disease, the distribution of causal variants in the genome and its dependence on other factors. However, as illustrated in this study, estimates of the genetic disease parameters using existing methods vary substantially conditional on, not only the approach and model used, but also the GWAS used as input. Developing a reliable predictor of the AUC trend will remain challenging until more robust estimators are developed. Despite these limitations, some of the current estimators can still be used to investigate the maximum achievable AUC. Altogether our analyses suggest that, assuming rare variants do carry a fair share of heritability, the genetic variance carried by these variants might be poorly captured by genotyped variants, so that until more large-scale whole-genome sequencing become available, the maximum AUC will remain bounded by the variant coverage of existing GWASs. In agreement with previous works^32^, the present study also support the use of imputed rare variants, even those with poor imputation quality, as an effective intermediate step to improve PRS predictive power.

For most analyses we assumed for simplicity that the causal variants are randomly distributed and we only modelled the relationship between minor allele frequency and effect size. This is clearly an over-simplification of the true underlying model, and as stated above, there is now strong evidence that causal variants distribution also varies across the linkage disequilibrium pattern and functional annotations^14,15,27,33,45,46^. The relationship between those factors and the causal effect has not been fully specified, and is expected to vary across outcomes. Instead of investigating specific models, we derived the relationship between the variant coverage and a vast set of functional annotations. We observed significant variability across annotations, with both positive and negative enrichments in variant coverage. However, except for CDS that show substantially poorer coverage, those differences did not have any qualitative impact on our results.

Our study has multiple limitations. First, our study only includes six diseases. We initially intended to include more outcomes, but for all other diseases we considered (e.g. chronic obstructive pulmonary function, Crohn’s disease, hypertension, etc) there was not enough available data to ensure a fair and objective analysis. Second, for real data heritability, we used twins study heritability as a proxy for the total additive genetic variance. We appreciate that these estimates might be overestimated because of shared environment and non-additive effects (e.g. gene-gene and gene-environment effects)^47^. Third, we assumed the genetic effects are homogeneous in the populations studied. Future improvement in predictive performances might depend on our ability to integrate potential heterogeneity in genetic effects conditional on demographic characteristics, basic health parameters, and lifestyle^4,48,49^. For example, there are increasing evidence that heritability might vary with age and sex^50,51^. Fourth, in our derivation of the maximum achievable AUC, we assumed a relatively simple model, focusing on the minor allele frequency-effect size relationship. Future works might incorporate more complex modelling including, in particular linkage disequilibrium (LD) dependencies. Note that assuming enrichment for causal variants in low LD regions will only worsen the gap between genotype-based AUC and imputed-based AUC. Fifth, heritability and AUC were derived using disease prevalence reported by the CDC. Prevalence clearly varies across population and using other estimates will impact the absolute values of both parameters, but not their relative values though. Sixth, we focus all our analyses based on the gold standard approach for PRS, that is, using univariate GWAS with the largest possible sample size. There is now increasing literature on multitrait approaches^52-54^, where the predictive power can be boosted by partly circumventing the challenge of assembling very large sample size.

## Methods

### GWAS data assembling

We collected publicly available genome-wide association study (GWAS) summary statistics from six outcomes: type 2 diabetes, coronary heart disease, breast cancer, Alzheimer disease, asthma, and body mass index (BMI) which was used to study obesity. The design of each study was assessed carefully to ensure it matches stringent criteria for inclusion in our analysis: i) all studies had to include a majority of individual of European ancestry; ii) studies with limited genetic coverage, such as exome-wide screening and *ad hoc* genotyping chips (e.g. Metabochip or Immunochip) without imputation were excluded; iii) phenotype definition had to be relatively homogeneous, although we kept some study with heterogeneity for comparison purposes (early onset for asthma, and Alzheimer disease proxy derived from parent status). Note that for some meta-analysis, we also had access to GWAS results of individual cohorts, which were used as additional data points. After this quality control filtering, a total of 29 GWAS summary statistics with the baseline required data (coded allele, sign statistics and *P* value) remained for analysis (**Table S2**). To complete our panel of GWAS we conducted five additional GWAS in the UK Biobank using unrelated participants of European ancestry, and cases sampled from the entire cohort: breast cancer (N_cases_=5,000; N_controls_=50,000); coronary artery disease (N_cases_=5,000; N_controls_=50,000); asthma (N_cases_=1,000; N_controls_=10,000); and BMI (N=20,000 and N=80,000). All 34 GWAS were harmonized and converted to hg38 using the liftover package^55^. As one of the ultimate goals of these data is to compare the predictive accuracy of PRS scores in the FinnGen cohort, we only kept variants available in that dataset.

### Estimation of prediction performances of polygenic risk score in real data

For each GWAS, we derived a vector of individual variants PRS weights *γ*=(*γ* _1_, *γ* _2_ *… γ* _*M*_), where *γ*_*i*_ is the weight of variant *i* and *M* is the number of variants available in the GWAS, using the LDpred2 approach^7^, a popular Bayesian method. We used the ‘auto’ option that automatically tune hyper-parameters including the PRS sparsity and the SNP heritability, and the recommended number (30) of Gibbs sampling chains. European descent participants from the UKB biobank were used as a reference panel for LD derivation, and we restricted the analysis to the GWAS variants overlapping with a set of 1,054,330 HapMap3 variants^42^. For each GWAS, variants with an estimated effective sample size smaller than 50% or larger than 110% of the expected maximum were further filtered out to avoid a miscalibration of the estimated regression coefficients^56^.

We use those weights to construct genetic risk scores in two test datasets (**Table S3**). The scores were derived as *PRS*=*γ*^*t*^ *X*, where *X* is a matrix of genotypes from the test dataset with coded alleles matching the coded alleles of the original GWAS. The first dataset included 16,000 unrelated participants of non-European ancestry from the UK Biobank, where ancestry (Ashkenazi, Iranian, Indian, Chinese, Caribbean, and Nigerian) was derived following Privé et al^18^. This analysis was also used for the fine tuning of the PRS and to derive the average weights over the 30 Gibbs sampling. The second dataset included 392,423 participants of Finns ancestry from the FinnGen cohort (see **Supplementary Notes**). Except if specified otherwise, we defined the effective sample size of a binary outcome GWAS as *N*_*eff*_ =4 / [1/ *N*_*case*_ +1/ *N*_*controls*_], which re-scales cohorts sample size with unequal numbers of cases and controls to the same unit^57^. We estimated the predictive performance of the PRSs using the Area Under the Roc Curve (AUC). The AUC is a commonly used classifier in genetics, which value equals to the probability of a randomly chosen case to be ranked higher than a randomly chosen control. For the non-European UK Biobank samples, we first derived the AUC per ancestry (**Table S3, Fig. S2**), and combined results using a standard inverse-variance meta-analysis: 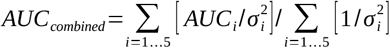 and 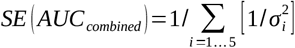 where and 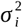 are the AUC and variance derived for population *i*=1 *…* 5.

### Variance captured by genotyped and imputed variants

Consider a standardized phenotype *Y* drawn from a polygenic model and defined as the linear additive effect of *M* standardized causal variants. Its variance equals *V*_y_ =*h*^2^+ *V*_e_, where *h*^2^ is the genetic variance, or heritability and *V* _*e*_ is the residual environmental variance.

Assuming the genetic effects at the causal variants are independent of variants’ correlation, the heritability can be approximated as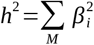, where *β*_*i*_ is the effect of a standardized variant *i*.

We derived the proportion of *h*^2^ that can be recovered given an infinite GWAS sample size, when only a subset *S* of variants have been genotyped, while the remaining *M ∉ S* are either imputed or missing. We considered two metrics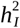 the genetic variance captured when using all *M* variants whatever their imputation quality, and 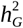 the genetic variance captured when using only the subset *S* of genotyped variants, but accounting for the effect of the *M ∉ S* variants captured through linkage disequilibrium. The first metric 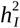is defined as 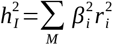 where 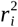 is the squared correlation between the sequence variant *i* and its imputed value. The second metric 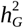 is defined as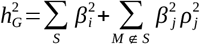 where 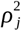 is the squared correlation between the untyped variants *j* and the genotyped ones, obtained from a multiple regression (see below for the derivation).

Both features were derived using a subset of 125,152 participants of European ancestry from the UK Biobank with genome-wide genotyping and whole genome sequencing data available. Note that for 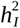, we first considered using the imputation quality score, commonly referred as the *info_score*, as defined by Marchini & Howie^58^, which is expected to be an unbiased estimator of *r*^2^. However, this metric appears to be inflated in both simulated and real data, especially for variants with low minor frequencies (**Fig. S8**). We therefore derived *r*^2^ using the aforementioned individual-level sequencing data. We used the imputed SNP-array data provided by the UK Biobank. We lifted-over the imputed variants for the 125,152 samples, retaining 99.5% of the original imputed sites in GRCh38. We then derived an r2 between the imputed dosages and the whole-genome sequencing data for each of the overlapping sites.

.The *r*^2^ was estimated from a standard univariate linear model using each sequenced variants as the outcome and the imputed ones as predictors. For 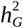, we used genotyped variants available on the *UK Biobank* Axiom array^21^ as our baseline. The *ρ*^2^ were derived as the adjusted squared-correlation obtained from a standard multiple linear regression with the *lm()* function from the R software. Each non-genotyped variant *j* was predicted from a set *Ω* of neighboring variants in a window of ±1.5Mb (**Fig. S9a**). For simplicity we derived *r*^2^ and *ρ*^2^ using only variants from chromosome 22, under the assumption that estimates from this chromosome are representative of the entire genome. There are 12,968 variants genotyped on this chromosome. Out of 659,092 variants available from the sequencing data, a total of 503,195 remained for analysis after removing those with a minor allele frequency (MAF)< 0.001%. As expected, the two metrics are highly correlated (*cor* (*ρ*^2^, *r*^2^)=0.71) but, the average *r*^2^ was substantially larger than 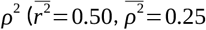 **Fig. S9b**).

In our simulations, the proportion of heritability captured was derived using both parameters (*r*^2^ and *ρ*^2^), and *β*=(*β*_1_*… β*_*M*_) coefficients drawn from a normal distribution using the alpha model^36^. The alpha model assumes an inverse relationship between the variant frequency and the per-allele effect, with rare causal variants harboring larger per-allele effect than common variants. This model has been empirically confirmed in many studies (e.g. ^59^). In practice, the genetic effect of variant *i* is drawn from the following:

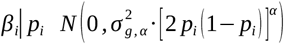 where *p*_*I*_ is the minor allele frequency, and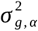 is a constant constraining the outcome heritability (**Fig. S10a**). In our analysis we considered *α* values in the range [-1.5 ; 0.0], and draw random 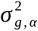 in [0,1]. We also investigated an attenuated alpha model where the contribution of the rarest variants was decreased using an *ad hoc* iterative weight function (**Supplementary Notes**).

### Maximum achievable AUC

The expected maximum achievable AUC is derived using the approximation proposed by Wray et al^3^: 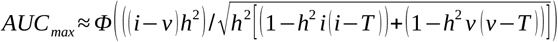where *h*^2^ is the heritability on the liability scale, *Φ* is the cumulative density function of the normal distribution, *z* is the height of the standard normal density at the threshold *T* =*Φ*^− 1^(1− *K*), and with *i*=*z* / *K* and *v*=− *z* / (1− *K*), where *K* is the disease prevalence. We confirmed the validity of the approximation using a simple simulation model involving independent causal variants with a linear additive effect on the outcome and using *h*^2^ in [0.2 ; 0.7] and the prevalence *K* in [0.01 ; 0.25] (**Fig. S4** and **Supplementary Notes**). Estimation of the AUC conditional on alpha (**Fig. 2b**) were derived by replacing *h*^2^ by either 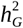 or 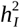, themselves derived based on alpha drawn in [-1.5, 0] and *β*=(*β*_1_*… β*_*M*_) coefficients generated from a normal distribution.

For real data analysis, disease prevalence from the six outcomes were pulled from the CDC website (https://www.cdc.gov/, **Table S6**). For the heritability, we considered various estimates : i) the total heritability derived from twins studies for coronary artery disease (0.55)^60^, type 2 diabetes (0.72)^61^, breast cancer (0.27)^62^, Alzheimer disease after excluding the effect of APOE (0.49)^63,64^, asthma (0.70)^65^, and body mass index (0.75)^66^ ; ii) 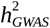, the heritability captured by GWAS variants derived using five alternative approaches: SBayesS ^24^, sumHer^25^, LDSC regression^26^, GENESIS^14^, and MiXeR^27^ (**Supplementary Notes**), and iii) 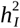, the heritability captured by all imputed variants. The later estimate requires three parameters: the total heritability, the proportion of heritability captured given alpha, and a value of alpha. For the total heritability, we used the twin study estimate. For the proportion of heritability captured, we used the estimate derived using the UK Biobank sequencing data. The choice of alpha was more challenging and is described below.

Most real data estimates of the GWAS variants heritability 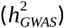 are derived based on genotyped variants along a modest subset of imputed variants with very high quality (typically an *info_score* ≥ 0.8). As a results, assuming our estimation of the heritability captured by genotyped variants based on the UK Biobank sequencing data (**Fig. 2a**) is valid, the previously described 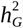 is expected to approximate 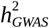 providing a relevant value of alpha. We assessed the equality between the two metrics using alpha derived from various approaches (SBayesS, sumHer, and individual-level data from the UK Biobank, **Table S6**), but none of them match the expected based on 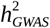. Therefore, for each disease, we selected a “*Best fit”* alpha so that 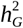 is equals the median of 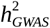 derived over the five aforementioned approaches (**Table S6**). This “*Best fit”* alpha was used to derived 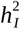 for the six outcomes.

### Genetic coverage and functional annotations

We investigated the association between *r* ², the squared-correlation between the true and the imputed genotypes and a total of 1,099 functional annotation pulled from nine sources: baseline GENCODE annotations: intron, gene, exon, CDS (coding DNA sequence), tss (transcription start site), tts (transcription termination site), and UTR (untranslated regions) ; and epigenetic annotations across a vast range of tissues and cell types: TFBS (transcription factor binding site) ; FANTOM5 (functional annotation of the mammalian genome version 5) ; promoters ; enhancers, and dyadic from Roadmap, DHS (DNase I hypersensitive sites) derived from two studies, and super enhancer. The analysis was conducted using the same data used to derive 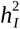 and 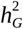 (503,195 variants from chromosome 22 from 125K UK Biobank participants of European ancestry).

### Data availability

All GWAS summary statistics have been downloaded from publicly available websites including dedicated page from consortia, the NHGRI-EBI Catalog of human genome-wide association studies, and the FinnGEN GWAS repository. Individual-level data from the UK Biobank were accessed from the UK Biobank Resource under Application Number 42260 and 66995. Individual-level data from FinnGEN were conducted by co-authors from the University of Helsinki with privileged access.

### URL resources

GWAS catalog: https://www.ebi.ac.uk/gwas/

GCTA: https://yanglab.westlake.edu.cn/software/gcta/

SBayesS: https://cnsgenomics.com/software/gctb/

MiXeR: https://github.com/precimed/mixer

GENESIS: https://github.com/yandorazhang/GENESIS

LDSC regression: https://github.com/bulik/ldsc

sumHer: https://dougspeed.com/sumher/

FinnGen results: https://risteys.finngen.fi/

HapMap3: https://www.sanger.ac.uk/resources/downloads/human/hapmap3.html

CDC disease prevalence: https://www.cdc.gov/datastatistics/index.html

Functional annotations: https://github.com/gkichaev/PAINTOR_V3.0/wiki/2b.-Overlapping-annotations

## Acknowledgments

We want to acknowledge the participants and investigators of the FinnGen study and the UK Biobank cohort. This research has been conducted using the UK Biobank Resource under Application Number 42260 and 66995. We also would like to thanks the authors of GENESIS, sumHer and SBayesS for their helpful recommendations.

This work has been conducted as part of tshe INCEPTION program (Investissement d’Avenir grant ANR-16-CONV-0005). This research was supported by the Agence Nationale pour la Recherche (ANR-20-CE36-0009-02 and ANR-20-CE15-0012-01).

## Authors contributions

HA and LH conceived and supervised the project. LH, JK, ZY, SR, GP carried out the primary analyses. CA, CB, HJ, AF, YS conducted secondary analyses. ZY and AG led the analyses involving individual-level data from FinnGEN. SR, OD and LH led the analysis involving individual-level sequencing data from the UK Biobank. FP, BJ, CA and HA designed the LDpred2 pipeline for PRS analysis. HA and LH co-wrote the manuscript with input from all other authors. All authors contributed to discussions.

