## Supplement for "Polygenic risk score prediction accuracy convergence"

### Supplementary Material

### Supplementary Notes

#### *Existing AUC from the literature*

We conducted in spring 2020 a *PubMed* search for studies reporting the prediction accuracy of genetic risk score for six outcomes, coronary artery disease (CAD), breast cancer (BRCA), type 2 diabetes (T2D), Alzheimer disease (AD), asthma, and obesity using the disease name and the terms "*AUC*" and "*genetic risk score*". We identified a total of 57, 48, 86, 25, 9 and 48 publications for CAD, T2D, BRCA, AD, asthma, and obesity, respectively, which we completed with a few additional studies identified using *ad hoc* search terms. We conducted a first screening and filtered out irrelevant studies not reporting AUC. For the remaining ones, we carefully checked the analysis pipeline, and kept studies using i) at least 300 cases (the number of controls was always larger than the number of cases), ii) using mostly participants of European ancestry, and iii) where the AUC was derived using the genetic risk score only, excluding other covariates (*e.g.,* age, sex, etc). After this careful quality control, a total of 6, 20, 8, 5, 3 and 7 studies remained for CAD, T2D, BRCA, AD, asthma, and obesity, respectively. For each study we extracted the following parameters: the number of cases and controls of the training set used to derive the polygenic risk score, the number of genetic variants used, the reported AUC in the validation set, and the publication year. All collected information are reported in **Table S1**. Trends in AUC increase as a function of sample size was tested using a weighted linear regression as implemented in the R *lm()* function, where weights were defined as the logarithm of the effective sample size ($N_{eff} = 4\,N_{cases} \times N_{controls}\,/(N_{cases} + N_{controls})$).

#### *Additional GWAS within UK Biobank*

Consortium-based GWAS provided results for large sample size for each disease considered in this study (**Table S2**). However, we found only limited publicly available GWAS results with intermediate sample size for multiple outcomes. To complete our prediction accuracy analyses, we conducted five additional GWAS in the UK Biobank using unrelated participants of European ancestry, and cases sampled from the entire cohort: breast cancer ($N_{cases}$=5,000; $N_{controls}$=50,000); coronary artery disease ($N_{cases}$=5,000; $N_{controls}$=50,000); asthma ($N_{cases}$=1,000; $N_{controls}$=10,000); and BMI (N=20,000 and N=80,000). We also conducted a GWAS for obesity from the whole sample, where obesity was defined as BMI>30 ($N_{cases}$= 74,660 ; $N_{controls}$=238,048) for comparison purposes. All analyses were conducted using Plink2.0's --glm option and adjusted for the top 15 principal components[1], age, and sex. The GWAS only included the ~1M variants from HapMap3 used in the construction of the PRS scores. For breast cancer, we selected female cases and controls and adjusted for 15 PCs and age. For each study, we conducted a stringent QC with a minor allele frequency (MAF) filter of 0.01, removing SNPs and individuals with missing data over 10%, and removed SNPs with Hardy-Weinberg equilibrium exact test p-value below 1e-50.

#### *Phenotype selection in UK Biobank and FinnGen*

For all analyses involving the UK Biobank and the FinnGen cohorts, we had to select a specific variable from all available fields for each of the six outcomes considered: type 2 diabetes (T2D), breast cancer (BRCA), coronary artery disease (CAD), Alzheimer disease (AD), asthma (AS), and body mass index (BMI)/obesity. In the FinnGen (https://risteys.finngen.fi/), we used the broadest definition of the disease or trait available. For type 2 diabetes, we used *Type 2 diabetes, definitions combined* (ID=T2D). For breast cancer, we used *Malignant neoplasm of breast* (ID=C3_BREAST). For CAD, we used *Major coronary heart disease event* (ID=I9_CHD). For Alzheimer, we used *Alzheimer disease* (ID=G6_ALZHEIMER). For asthma, we used *Asthma* (ID=J10_ASTHMA). For obesity, we used *Obesity* (ID=E4_OBESITY). In the UK Biobank (https://biobank.ndph.ox.ac.uk/), phenotypes were defined based on clinical ICD10 diagnosis codes (variable ID=41270) for five outcomes: breast cancer (ICD10 code=C50), coronary artery diseases (ICD10 codes=I21-I25), obesity (ICD10 code=E66), T2D (ICD10 codes=E10-E14), and asthma (ICD10 codes=E10-E14). For Alzheimer disease, we used AD-by-proxy (code 10 for variable ID=20110 from the mother and variable ID=20107 from the father), as proposed by Janssen et al[2], because the sample size for the Alzheimer disease status was too small. Counts of cases and controls for each outcome and each population considered are provided in **Table S3.**

#### *Correlation between European and FinGenn GWAS*

The GWASs we used to derive the PRS include a vast majority of individuals of European descent (**Table S2**). Although the Finns population from the FinGenn cohort are also of European descent, they have a slight north Asian admixture[3]. To explore possible PRS portability issues[4], we compared top association results from the largest GWAS[5-8] from **Table S2** with those from the FinnGen GWAS, publicly available on their website (https://www.finngen.fi/en/access_results). For the latter, we used summary statistics from release 5, using IDs listed in the previous section. For each outcome we extracted from the corresponding European and Finns GWAS pair, all SNPs with a *p*-value below 1e-6 in either study. We created a merged list of variants and kept only those for which a *p*-value was available in both GWAS. We next clumped this subset using the *ld_clump()* function from the R package "ieugwasr" (https://rdrr.io/github/MRCIEU/ieugwasr/man/ld_clump.html) with an $R^2$ threshold of 0.05 and default options, and the 1000 genomes[9] as a reference panel. Using either the European *p*-values or the FinnGen *p*-values for clumping produced qualitatively similar results. Overall, when plotting the estimated regression coefficient from each pair of studies against each other, we found highly consistent results between FinnGen and European GWAS (**Figure S1**).

#### *Specificity of the genetic model for obesity*

Along the study we use body mass index (BMI) GWAS to build polygenic risk scores and estimate disease parameters for obesity (defined as BMI ≥ 30 kg/m2), because the sample size available for this phenotype is substantially higher than for GWAS of obesity. In theory, the heritability of BMI should equal the heritability of obesity on the liability scale, and the disease parameters should be fairly similar. Indeed, obesity follows the definition of the liability threshold model (LTM) that is commonly used to report the heritability of binary outcomes[10], with BMI being the liability. Briefly, the liability describes the combined risk of genetic and environmental factors that contribute to the development of a disease, with the disease status being determined by the liability reaching a threshold. We did not conduct a comparison between the two outcomes because of the lack of data for obesity, however, we did notice differences in some parameter estimations when using either BMI or obesity (e.g. **Fig. S11**), that might be investigated in future studies.

Note that for the derivation of the predicted AUC of obesity using GENESIS (see next section), we used the BMI GWAS, while the approach is expecting z-score derived from a logistic regression. We investigated the possible impact of using BMI GWAS as a proxy for obesity GWAS through simulations. We generated series of replicates where a normally distributed outcome $Y$ depends on a single predictor $X$, and applied for each replicate a linear model $Y \sim \beta_0 + \beta_Y X$ and a logistic regression $logit(\Pr(D)) \sim \alpha + \beta_D X$, where $D$ is a dichotomized version of $Y$, defined as $D = 1$ if $Y$ is larger than the 70th percentile of $Y$ and $D = 0$ otherwise (mimicking the BMI-obesity relationship). We compared the z-scores from the two models $z_Y = \hat{\beta}_Y / \sigma_{\hat{\beta}_Y}$ and $z_D = \hat{\beta}_D / \sigma_{\hat{\beta}_D}$ and found strong correlation ($r^2 > 0.9$ across the scenarios considered), but with $z_D$ being systematically smaller than $z_Y$, with on average $z_D = 0.73\, z_Y$, corresponding approximately to a two-fold decrease in sample size for the dichotomized outcome. We used this *ad hoc* correction factors when plotting the obesity GENESIS results (**Fig. S5**).

#### *Specificity of the genetic model for Alzheimer disease*

Genetic variants within the Apolipoprotein E (ApoE) gene encode for three haplotypes, ε2, ε3 and ε4 that are critical determinants of Alzheimer disease risk[11]. Those variants with large effect deviate from the standard polygenic model and are commonly assessed separately from the effect of other variants. The causal APOE variants were available in the three largest Alzheimer disease GWAS (Lambert et al. 2013, Kunkle et al. 2019 and Jansen et al. 2019), but absent from the smallest one (Li et al. 2008), thus potentially impacting the comparison of the predictive performance. Moreover, the LDpred2[12] approach used to derive the polygenic risk score (PRS) can be sensitive to such deviation of the polygenic model. To address this possible limitation, in complement to the standard pipeline, we derived three additional predictive models: a PRS excluding the APOE region (PRS.noAPOE), a risk score based on APOE ε2 and ε4 allele genotypes (rs429358 and rs7412) with effect estimates pulled from Kunkle et al[13] (the largest GWAS for AD status) (PRS.APOE), and a PRS including PRS.noAPOE and PRS.APOE as two distinct components. The predictive performances from these alternative PRSs are presented in **Figure S3**. Also, note that when using the PRS without APOE in **Figure 2**, we used the twin heritability estimates after subtracting the contribution of APOE variants, which has been reported to be approximately 9% of the total phenotypic variance[14].

Another challenge in the analysis of Alzheimer disease is the use of the so-called AD-by-proxy instead of the AD status in the largest GWAS (Jansen et al[2]). AD-by-proxy is a score determined by the Alzheimer status of the parents' participants. It has been proposed to virtually increase the available sample size in AD GWAS and therefore increase statistical power. In our estimation of the AUC in the FinnGEN cohort, we use the true AD status. This might potentially explain the limited increase in AUC in that cohort when using the Jansen et al PRS. Conversely, in the estimation of the AUC in the non-European UKB participants, we used the AD-by-proxy as the primary outcome because of a limited sample size for the true AD status (N=961 for AD-by-proxy cases, and N=65 for AD case). For that cohort, the AUC show a substantial increase for the Jansen et al PRS, likely due to the outcome correspondence. While a full investigation of the impact of using AD-b-proxy is out of the scope of this study, we derived the predictive power of the APOE variants (PRS.APOE) in the six non-European UK Biobank populations as a qualitative marker of this proxy phenotype. We obtained AUC=0.65 (SD=0.58) and AUC=0.54 (SD=0.52), for the AD status and the AD-by-proxy status, respectively. This is in line with the expectation that a loosen disease definition can impact the predictive power of genetic risk score.

#### ***Parameters estimation from GWAS using existing tools***

We used multiple tools to estimate disease genetic parameters from GWAS summary statistics. We applied GENESIS[15], SBayesS[16], sumHer[17], LDSC[18], and MiXeR[19,20] to all GWAS from **Table S2** except for Saxena et al 2007 and Li et al 2008 because of modest sample size ($N_{eff}$=2,931 and 1,489, respectively) (**Table S5** and **Fig. S5-S7**). All methods estimate heritability. GENESIS, SBayesS, and MiXeR additionally estimate the number of causal variant and the polygenicity. SBayesS and sumHer also provide estimates of $\alpha$, the MAF-effect size relationship parameter. GENESIS output was also used to derive the expected trend of the AUC as a function of sample size (**Fig. S5**). We parametrized the method following the available tutorials and direct recommendations from the authors. Note that those methods sometimes refer to parameters that have different definition or labels (e.g. $\alpha$ is sometimes referred to as *S*, effective sample size is defined differently across methods, the output heritability scale is heterogeneous, etc). When possible, we harmonized the output estimates to allow for a direct comparison across methods. Below is a brief description of the overall analysis pipeline and specific parametrization of each method.

All analyses were conducted using GWAS variants overlapping with a reference set of 1,054,330 variants from HapMap3[21]. Most of these methods assume the original GWAS have not been corrected for chi-squared inflation using genomic control (GC)[22]. We carefully read each GWAS paper to assess whether GC correction was applied. The correction strategy across GWASs was heterogeneous (**Table S2**), sometimes applied at the study level (by each study, before the meta-analysis), or at the meta-analysis stage, at both, or neither. The information was also sometimes missing, or partial (a correction was reported, but the actual correction factor was not provided). We identified three studies were the inflation factor $\lambda_{GC}$ use for correction was fairly large (Nikpay et al. 2015, Speliotes et al. 2010, and Locke et al. 2015) and we back-corrected the GWAS results before the analysis.

GENESIS produces estimates of the SNP heritability, the number of causal variants, and allows to generate expected AUC as a function of the sample size. Following recommendations from the authors, we ran a three-component mixture model, which assumes that the effect sizes for susceptibility variants can be described by two distinct normal distributions (small and large effects) and a third component corresponding to non-associated variants. We use as input the effective sample size derived as $N_{eff} = N_{cases} \times N_{controls} \,/\, (N_{cases} + N_{controls})$, but report the results using the alternative derivation $N_{eff} = 4\, N_{cases} \times N_{controls} \,/$

$(N_{cases} + N_{controls})$. The heritability estimated in GENESIS is on the log-odds scale and was transformed to the liability scale using the formula $h_l^2 = h_{log}^2 K^2(1-K)^2/z^2$, where $K$ is the disease prevalence in the population and $z$ the height of the standard normal probability density function at the liability threshold[10].

SBayesS produce estimates of the number of causal variants, the SNP heritability and $\alpha$. We used the shrunk LD matrices proposed along the software. It requires the frequency of the coded allele, which was not available for several GWAS. Instead, we used frequencies derived from participants of European ancestry in the 1KG reference panel. For all analyses we used the options *--exclude-mhc* to remove the MHC region from the derivation, and *--impute-n* that re-derive the sample size, as the per-variant sample size is likely heterogeneous for many of the GWAS analysed. This option also filters out variants that have sample size 3 standard deviation away from the expected. For the sample size, we used the total sample: $N = N_{cases} + N_{controls}$. Heritability on the liability scale was derived in a second step using the formula[10]: $h_l^2 = h_{obs}^2 \left(K(1-K)\right)^2/(P(1-P)z^2)$, where $K$ and $P$ are the population and in-sample prevalence, respectively.

MiXeR[19,20] produces estimates of the fraction and number of causal SNPs and the SNP heritability, based on GWAS summary statistics and assuming a Gaussian mixture model. MiXeR takes into account SNP heterozygosity, LD structure, and residual inflation of z-scores due to variance distortion (which can arise from cryptic relatedness in the sample). We used the default reference panel, the 1000 Genomes Phase3 data and the MHC region was excluded from the analysis. As recommended by the authors, we used the standard definition of the effective sample size: $N_{eff} = 4\, N_{cases} \times N_{controls} / (N_{cases} + N_{controls})$. Heritability on the liability scale was derived in a second step using the formula[10]: $h_l^2 = h_{obs}^2 \left(K(1-K)\right)^2/(P(1-P)z^2)$, where $K$ and $P$ are the population and in-sample prevalence, respectively.

LDSC regression[18] was used to derive estimates of SNP heritability. Here we used the original model proposed in 2015 by Bulik-Sulivan et al[18], which has been widely used in the field. Heritability is derived from a regression between the Chi-squared for association and the LDscore, a metric that quantifies the strength of the correlation between each variant and its neighbours. The standard model implicitly assumes an infinitesimal model with an equal per-variant contribution to the heritability. For the sample size, we used the total sample: $N = N_{cases} + N_{controls}$. Heritability on the liability scale was derived in a second step using the formula[10]: $h_l^2 = h_{obs}^2 \left(K(1-K)\right)^2/(P(1-P)z^2)$, where $K$ and $P$ are the population and in-sample prevalence, respectively.

sumHer[17] is an extension of the LDAK model[23] for summary statistics. As compared to LDSC, sumHer uses a model were per-SNP heritability varies with both linkage disequilibrium (LD) and minor allele frequency (MAF). We used the options *--cutoff* 0.01 to remove variants that explain more than 1% of phenotypic variance, and specified the in-sample and population prevalence with *--prevalence* and *--ascertainment*, so that the heritability is directly derived on the liability scale. For the sample size we used $N = N_{cases} + N_{controls}$. For the estimation of $\alpha$ we used the solution proposed within sumHer that consists in testing a range of $\alpha$, and estimating the best fit (i.e. the highest log-Likelihood) using the *--find-gaussian* option.

### *Simulations to study the expected convergence of PRS*

To characterize the convergence of PRS prediction as a function of the disease parameters and sample size, we simulated series of matched case/control (i.e. $n = n_{case} = n_{control}$) replicates varying the disease prevalence ($K$ = [0.01; 0.25], the disease heritability ($h^2$ = [0.2; 0.7], the number of causal genetic variants ($M$ = [100, 1000, 5000, 10,000], and the sample size used in the GWAS ($n$ = [250; 2,500; 25,000; 250,000]). Independent causal single nucleotide polymorphisms (SNP) were generated under a binomial distribution with minor allele frequency randomly drawn in [0.05, 0.5]. The disease status was generated under a liability threshold model. The effect of the SNPs on the liability were drawn from a gaussian distribution $\beta \sim \mathcal{N}(0, h^2/M)$, and the threshold defined as $\mathrm{T} = \Phi^{-1}(1-k)$ where $\Phi$ is the cumulative distribution function of the normal distribution. Each replicate was split in two subsets. The first subset was used to estimate genetic effect $\hat{\beta}$ for each of the $G_{i=1\ldots M}$ variants using univariate logistic regression. A PRS was then computed in the second subset as: $PRS = \sum_{i=1\ldots M} \hat{\beta}_i\, G_i$. The predictive performances of the PRS for each model was derived as the average over 10 simulations of the Area under the ROC (AUC) curve in this second subset.

#### *Using the* info_score *as a proxy for the imputation squared correlation*

We initially considered using the imputation quality score, commonly referred as the *info_score,* as defined by Marchini & Howie[24], as a proxy for $r^2$, the squared-correlation between true and imputed variants. To assess the validity of this assumption, we conducted a simulation study to compare the *info_score* metric, as commonly derived using the IMPUTE suite[24], with the $r^2$ derived from a linear regression when using the same information. We extracted a dataset of $N$=20,000 participants and 1,000 neighbouring variants from the imputed UK biobank data with *info_score* larger than 0.3, and MAF larger than 1%. For imputed variants, we used best guess genotypes. We split the dataset into a training group (n=10K) and a testing group (n=10K), and hid a random subset $H$ of 900 variants in the training group to be imputed using the remaining subset $G$ of 100 variants. However, we excluded from the later set all variants with an absolute correlation above 0.7 with any other variant, resulting in a total of 35 variants used for the imputation. For each hidden variant $H_i$ with a minor allele frequency above 0.05 (n=822), we derived the squared-correlation $r^2$ and the *info_score* $I^2$*.* When all coefficients are standardized, the squared-correlation is $r^2 = \hat{\beta}^t \Sigma^{-1} \hat{\beta}$ where $\hat{\beta} = \hat{\beta}_1 \dots \hat{\beta}_{35}$ and $\Sigma$ are the estimates and covariance matrix from a standard multiple regression model defined as: $H_i\beta_0 + \sum_{t=1\dots 35} \beta_t G_t$. The *info_score* is derived as a function of $p_{ij1}$and $p_{ij2}$, the imputed probabilities for genotype $H_i = 1$ and $H_i = 2$ for individual $j$, and $\hat{\theta}_i$, the allele frequency estimate of $H_i$. It is defined as $I^2 = 1 - \sum_{j=1\dots N}\left(f_{ij} - e_{ij}^2\right)/\left(2N\hat{\theta}_i\left(1-\hat{\theta}_i\right)\right)$, where, $e_{ij} = p_{ij1} + 2p_{ij2}$ and $f_{ij} = p_{ij1} + 4p_{ij2}$. To derived the imputed probabilities $p_{ij1}$and $p_{ij2}$, we ran a multinomial regression model using R package *mlogit*[25] in the training group. As showed in **Figure S8b**, the two metrics display a very high correlation (squared correlation > 0.95) for common variants (with a minor allele frequency above 0.05), but a substantial inflation of the *info_score* for variants with low frequency. Because of this limitation, and thanks to the recent release of whole-genome sequencing data from the UK Biobank, we discarded the use of the *info_score* and use instead $r^2$ derived from real sequencing data. Comparison of $r^2$ and *info_score* in the UK Biobank sequencing data confirm the overestimation of *info_score* in these data (**Figure S8a**).

#### *Estimation of alpha for common diseases using UK Biobank individual-level data*

We derived the $\alpha$ parameter for the six outcomes using individual-level genetic and phenotypic data from the UK Biobank using the approach described by Schoech et al[26]. In brief, we derived $\mathbf{A}_\alpha$, the genetic correlation matrices (GRM) conditional on a pre-specific $\alpha$ values in [-1, 0], and defined as $\mathbf{A}_\alpha = \mathbf{X}\mathbf{D}_\alpha\mathbf{X}^t$, where $\mathbf{X}$ is the genotype matrix and $\mathbf{D}_\alpha$ is a diagonal matrix with element $D_{ii} = [2p_i(1-p_i)]^\alpha$. For each outcome $Y$, profile likelihoods $L(\alpha)$ of the multivariate normal model $Y \sim \mathcal{N}\left(0, \mathbf{A}_\alpha\sigma_{g,\alpha}^2 + \mathbf{I}\sigma_\varepsilon^2\right)$, where $\sigma_\varepsilon^2$ is the residual variance, were derived for a range of $\alpha$ value using the GCTA software[27]. The final $\hat{\alpha}$ estimate was derived as $\hat{\alpha} = argmax(L(\alpha))$. Estimation was done using up to 261,028 unrelated self-reported "white British" participants with genetic data for 17,211,988 variants with minor allele frequency (MAF) equal or larger than 0.01%, and *info_score* larger than 0.8.

In practice, the estimation of this parameter from individual-level data poses severe computational challenges, and the GRM-MAF-LD package proposed by Schoech et al[26] was not runnable in these data. We followed the same procedure, but re-implemented each step and optimized the parameters for computational purposes. For the derivation of $\mathbf{A}_\alpha$, we modified the GRM function from PLINK 2.0[28], to include the $\alpha$ weighting. Due to memory constraints, GCTA could not be run using a GRM of several hundred thousand individuals at once. To address this limitation, we divided the dataset into eight equally sized random sub-samples, and estimated $L(\alpha)_s$ for each subsample $s$ using GCTA. The overall $L(\alpha)$ was then derived as the sum of the resulting log likelihoods: $L(\alpha) = \sum_{s=1\dots 8} L(\alpha)_s$. We considered a range of $\alpha$ values in [-1.3 ; 0], with 0.1 steps. Data points were then interpolated using a Locally Weighted Least Squares Regression as implemented in the R *loess()* function to obtain the final estimate of $\hat{\alpha}$. We validated the procedure using real genotype data from the UK Biobank cohort and simulated phenotypes (**Fig. S12a**). We also attempted to estimate alpha across the MAF strata, using three bins: [0.5, 0.1], [0.1, 0.01], and [0.01, 0.001]. However, this estimation was unreliable, displaying slow convergence because of the limited variance within each MAF bin (**Fig. S12b**).

#### *Attenuated alpha model*

The alpha model is defined as $\beta_i|p_i \sim \mathcal{N}\left(0, \sigma_{g,\alpha}^2 \cdot [2p_i(1-p_i)]^\alpha\right)$, where $p_i$ is the minor allele frequency of variant $i$, and $\sigma_{g,\alpha}^2$ is a constant constraining the outcome heritability. For small value of alpha, it implies the per-allele effect of rare variants can be several orders of magnitude larger than for common variants (**Fig. S10a**). As noted in Schoech et al[26], the fit of the alpha model in evolutionary forward simulations is expected to hold above a fairly low MAF threshold (MAF>0.6% in the reported example), but might overestimate the effect of variants with MAF below that threshold. Let's denote $\theta$ the scaling factor of effect size coefficient, so that $\theta_i = [2p_i(1-p_i)]^\alpha$. To account for possible deviations of the alpha model, we considered attenuated effect size $\theta_i^*$ for very rare variants. We defined the attenuation using an inverse logit function with a tipping point at the aforementioned MAF threshold $T$=0.6%: $\pi = 1/[1 + exp(\log(MAF) - \log(T))]$. The attenuation was applied recursively to the beta coefficient ordered by decreasing MAF: $\theta_i^* = \theta_{i-1}^* + (\theta_i - \theta_{i-1})(1 - \pi w)$, where $w$ is a weight in [0, 1] to set the strength of the attenuation. **Figure S10b** illustrates the effect of the proposed attenuation on the MAF-effect size relationship for $\alpha = -0.3$. We varied the weight $w$ over [0.1, 0.3, 0.5, 0.7, 0.85, 0.95, 0.99] and repeated the experiment from **Figure 2a**, deriving the expected proportion of heritability captured across this range of attenuated alpha model using either $r^2$, the squared correlation between the sequence and imputed variants (**Fig. S10c**), or $\rho^2$, the squared correlation between the untyped variants $j$ and the genotyped ones (**Fig. S10d**).

### Supplementary Figures

#### *Figure S1. European and FinnGen GWAS*

Beta coefficients at top associated independent variants from FinnGen GWASs and European ancestry GWASs for six outcomes: obesity (a), breast cancer (b), asthma (c), type 2 diabetes (d), Alzheimer disease (e), and coronary artery disease (f). Across all variants, squared-correlation equal 0.43 (obesity), 0.56 (breast cancer), 0.50 (asthma), 0.67 (type 2 diabetes), 0.55 (Alzheimer disease), and 0.66 (coronary artery disease). Note that for obesity the European GWAS results were extracted from a BMI GWAS, explaining the scaling difference in regression coefficient. For Alzheimer disease (AD), the European GWAS was derived using AD-by-proxy, inducing a larger variance in the outcome, and again difference in the scaling of the regression coefficient.

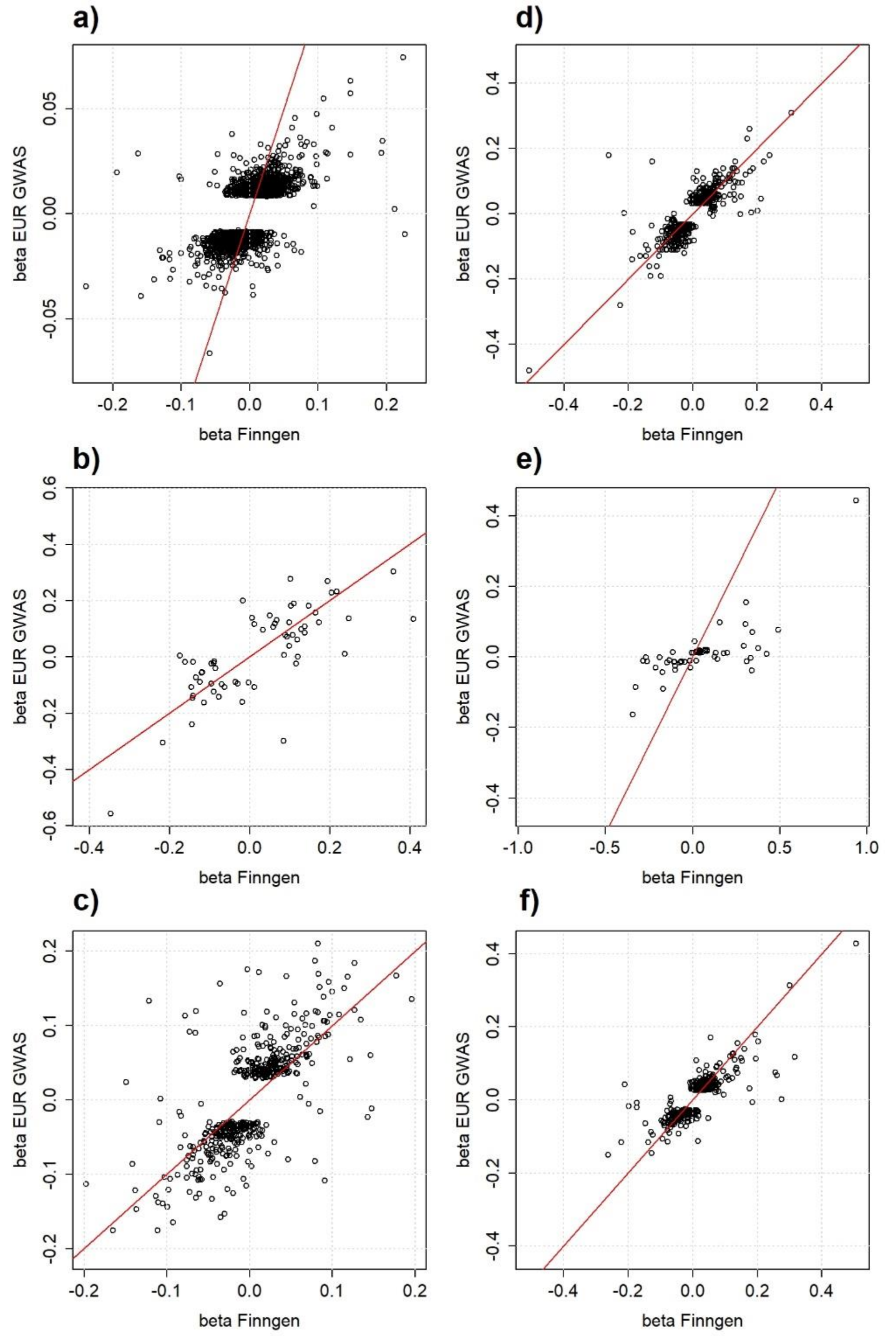

***Figure S2. AUC as a function of sample size for 6 UK Biobank populations***

AUC as a function of the effective sample size for each of the 6 ancestries analysed in the *UK Biobank* (Ashkenazi, Iranian, Indian, Chinese, Caribbean, and Nigerian). Grey areas correspond to the 95% confidence interval. AUC was derived for 6 outcomes: coronary artery disease (CAD), type 2 diabetes (T2D), breast cancer (BRCA), Alzheimer disease (AD), asthma, and obesity. Obesity was predicted using BMI GWASs. For AD, the outcome predicted was AD-by-proxy because of a limited number of AD cases within each ancestry. The PRS for this outcome was derived after excluding the APOE region.

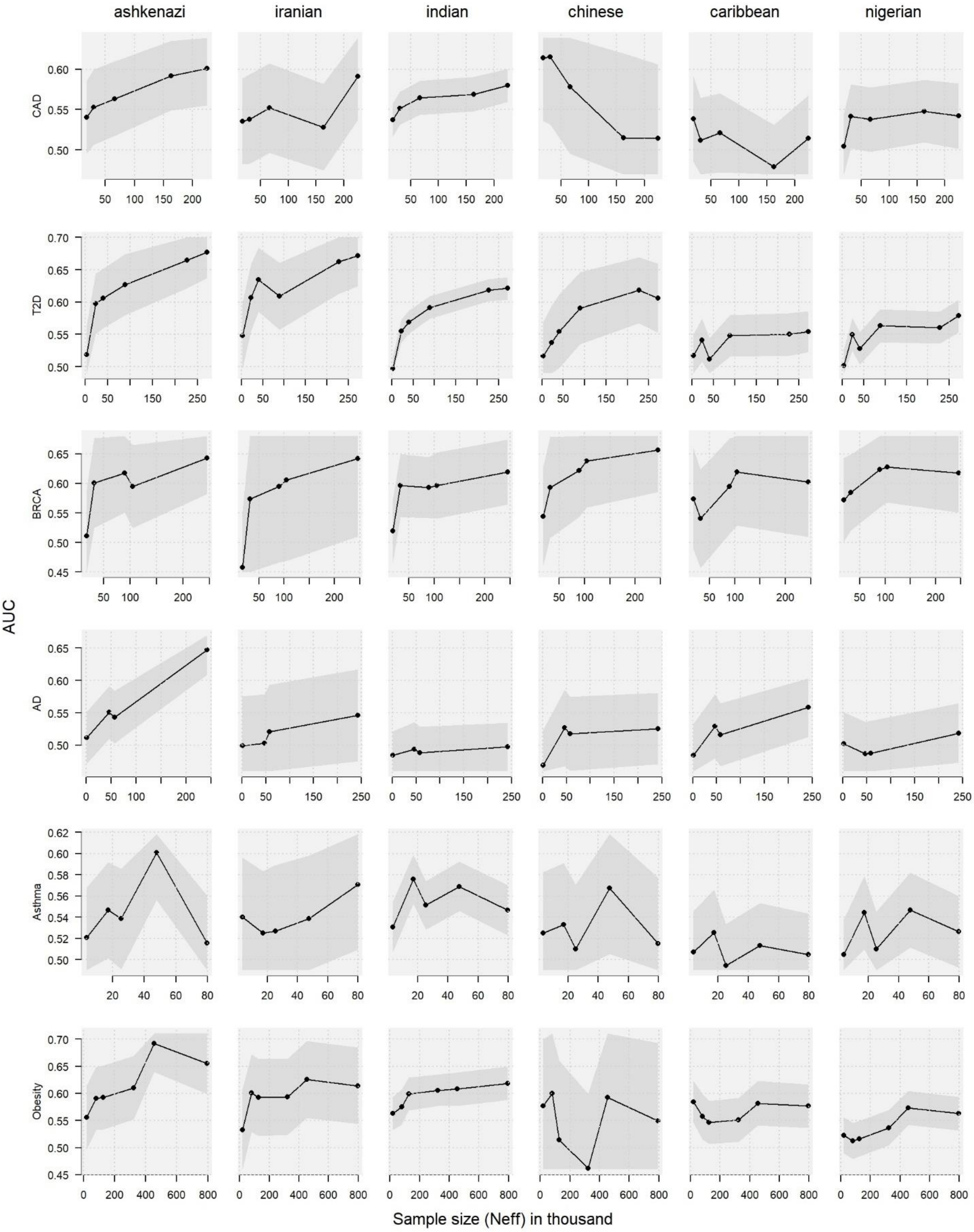

***Figure S3. Variability in Alzheimer prediction conditional on APOE***

For Alzheimer disease we considered three alternative genetic models conditional on the APOE region: i) a PRS derived after excluding the APOE region ("PRS excluding APOE", plain line), which was used as the main analysis, ii) a PRS derived using the same pipeline as for other diseases treating the APOE region as the rest of the genome ("*PRS including APOE region*", dash line), and iii) the combination of a PRS derived excluding the APOE region and score for the $\varepsilon_1$ and $\varepsilon_2$ variants as a distinct predictors ("*PRS and APOE treated separately and merged*", dotted line). The panels present the results of each of the three genetic scores in the FinnGen participants, and the non-European UK biobank participants per population and after a meta-analysis ("*UKB Combined*").

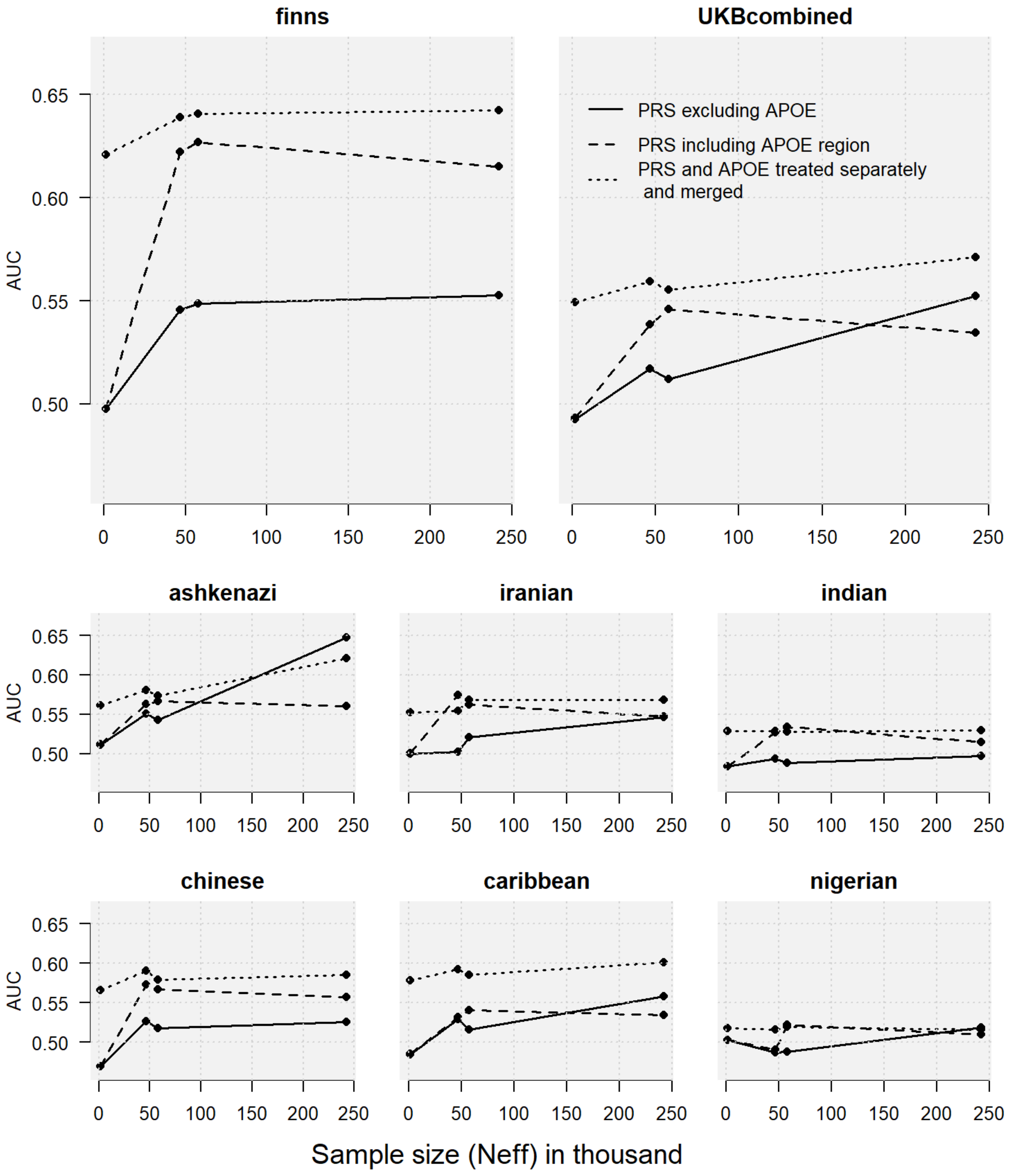

***Figure S4. AUC convergence using simulated causal variants***

Average predictive power as measured by the Area under the ROC (AUC) in simulated matched case/control replicates. Disease status was drawn under a liability threshold model using 100 to 10K independent causal variants. For each replicate, the effect of causal variants was estimated using univariate logistic regression using an increasing sample size ($n$ = [250; 2,500; 25,000; 250,000]). A PRS was then computed in an independent subset using those estimates and its performance derived as the average AUC over 10 simulations. Results across low and high heritability ($h_A^2$) and disease prevalence (K) are presented in panel a) ($h_A^2$=0.2, K=0.25), b) ($h_A^2$=0.2, K=0.01), c) ($h_A^2$=0.7, K=0.25), and d) ($h_A^2$=0.7, K=0.01). The black dash line represents the expected maximum achievable AUC derived based on heritability and prevalence.

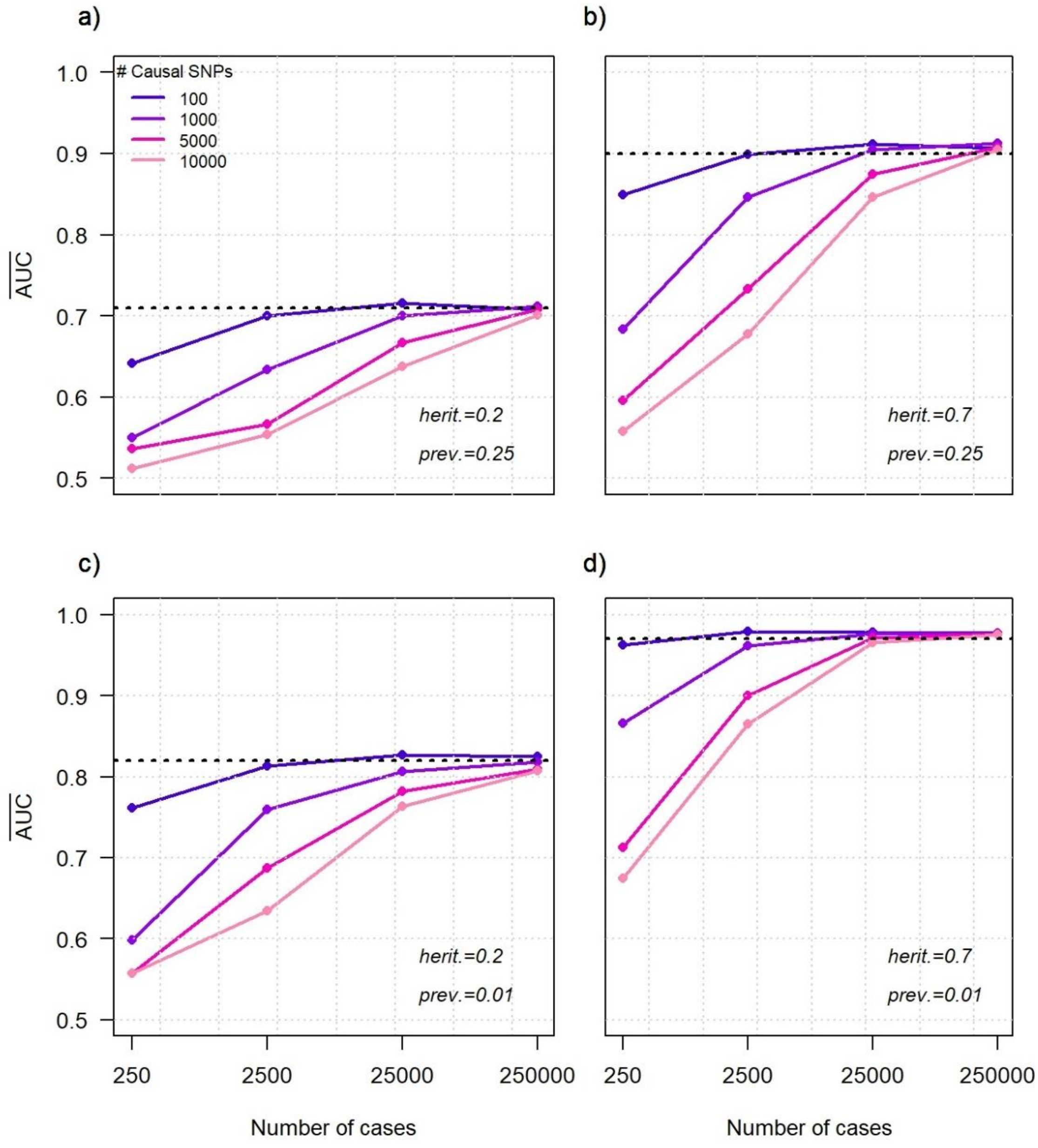

***Figure S5. AUC increase, heritability and number of causal variants derived from GENESIS***

Predicted AUC as a function of sample size derived from the GENESIS package assuming a 3-components genetic model for each of the six outcomes, coronary artery disease, breast cancer, type 2 diabetes, Alzheimer disease, asthma, and obesity, while using the available GWAS data. Note that for obesity, we used body mass index (BMI) GWAS and applied an *ad hoc* correction of the sample size, expected to match power of a binarized version of BMI. AUC derived in FinnGEN and the UK Biobank are indicated in black and red, respectively, for comparison (a). The predicted AUC are based on estimates of heritability (b) and the number of causal variants (c) estimated within GENESIS. Error bars indicate the 95% confidence interval. Note that for obesity predicted AUC, we used an *ad hoc* correction of the sample size to reflect the use of BMI GWAS instead of obesity GWAS.

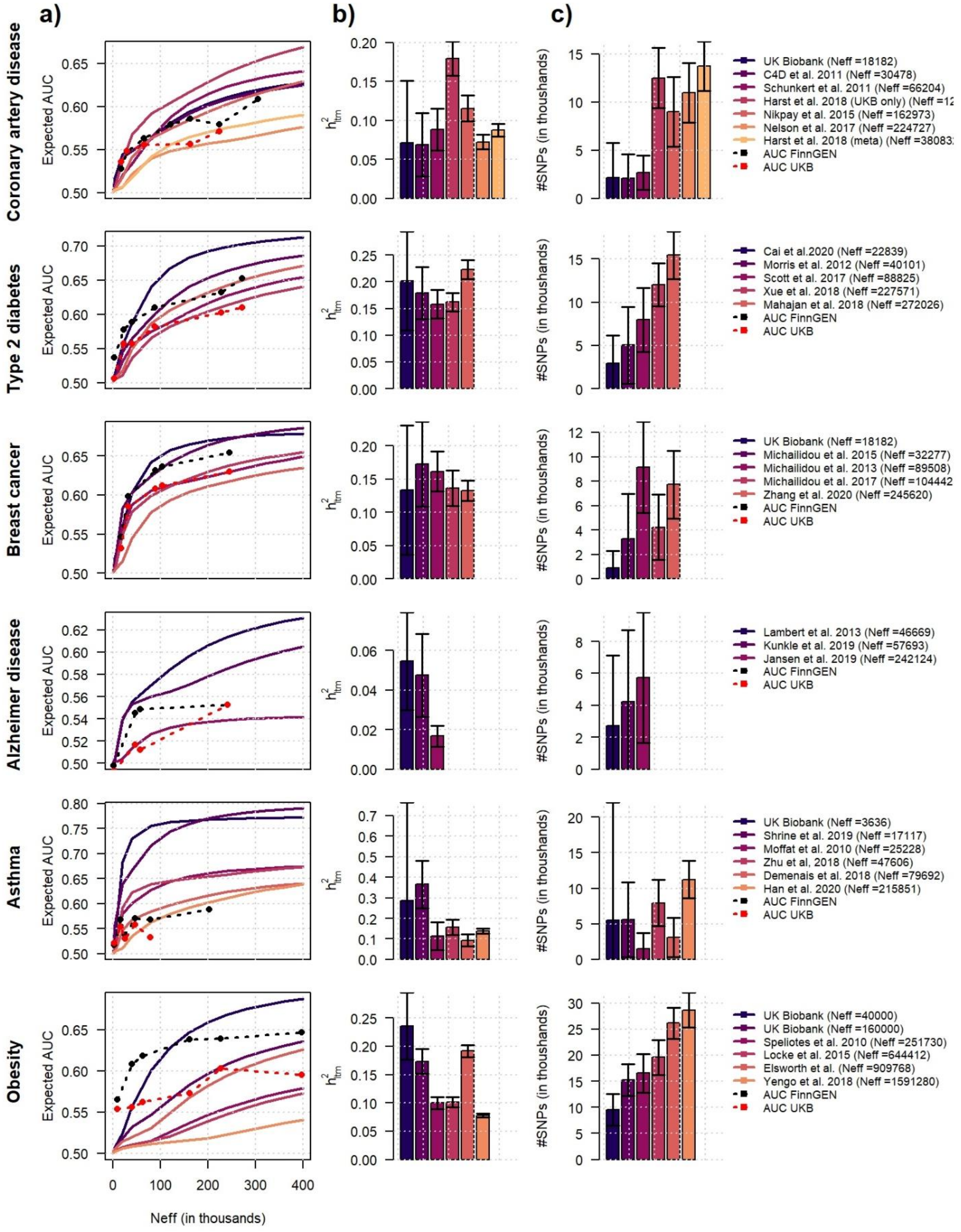

***Figure S6. Assessing the impact of GWAS heterogeneity on disease parameter estimation and AUC***

We conducted GWAS for body mass index (BMI) in random subsamples of the UK biobank with sample size $N$ increasing from 20K to 300K. For each GWAS, we applied SBayesS to estimate polygenicity (the percentage of causal variants), alpha, and heritability under the liability threshold model. For each resulting GWAS, we applied LDpred2 to derived PRS following the same pipeline as for Figure 1, and applied these PRSs to derive the AUC in a holdout test dataset from the UK Biobank not used in the GWAS. We repeated the analysis five times with a different holdout subset for each analysis. Panel a), b) and c) present estimates of polygenicity, alpha and heritability across the five experiments as a function of sample size. Panel d) presents the average AUC derived in the UK Biobank holdout (blue). Results AUC from Figure 1b (obesity), derived in FinnGEN ("Finns external") and in non-European participants from UKB ("UKB external") using PRS from external GWAS are presented in red and black, respectively.

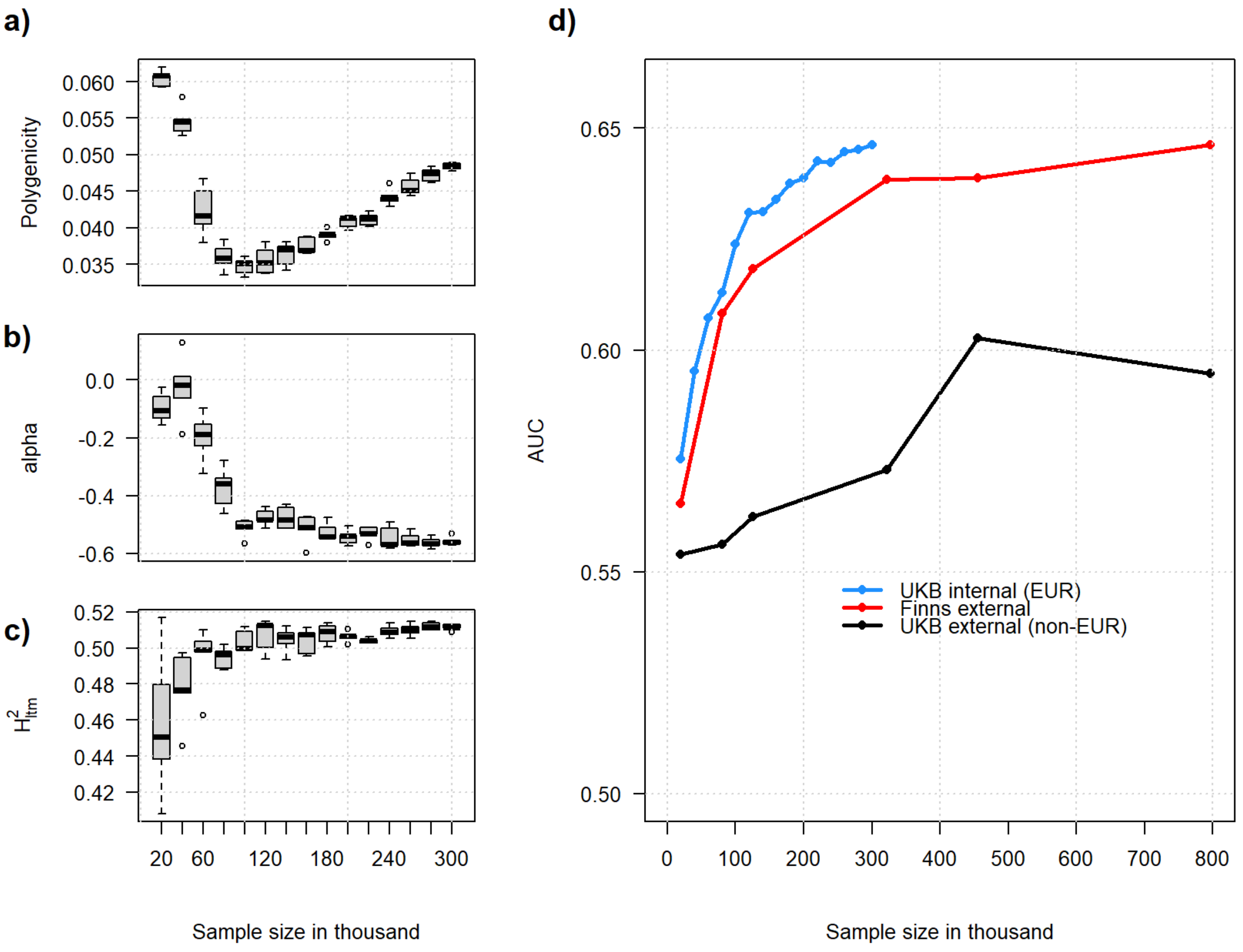

***Figure S7. SBayesS disease parameters estimates across the six outcomes and GWAS***

Estimates and 95% confidence interval of polygenicity (the proportion of causal variants), heritability and alpha derived using SBayesS across 31 GWAS summary statistics from six outcomes: coronary artery disease (CAD), breast cancer (BRCA), type 2 diabetes (T2D), Alzheimer disease (AD), asthma (AS), and body mass index (BMI). We only include GWAS with an effective sample size larger than 5,000. All analysis were conducted using a subset of approximately 1M hapmap3 variants.

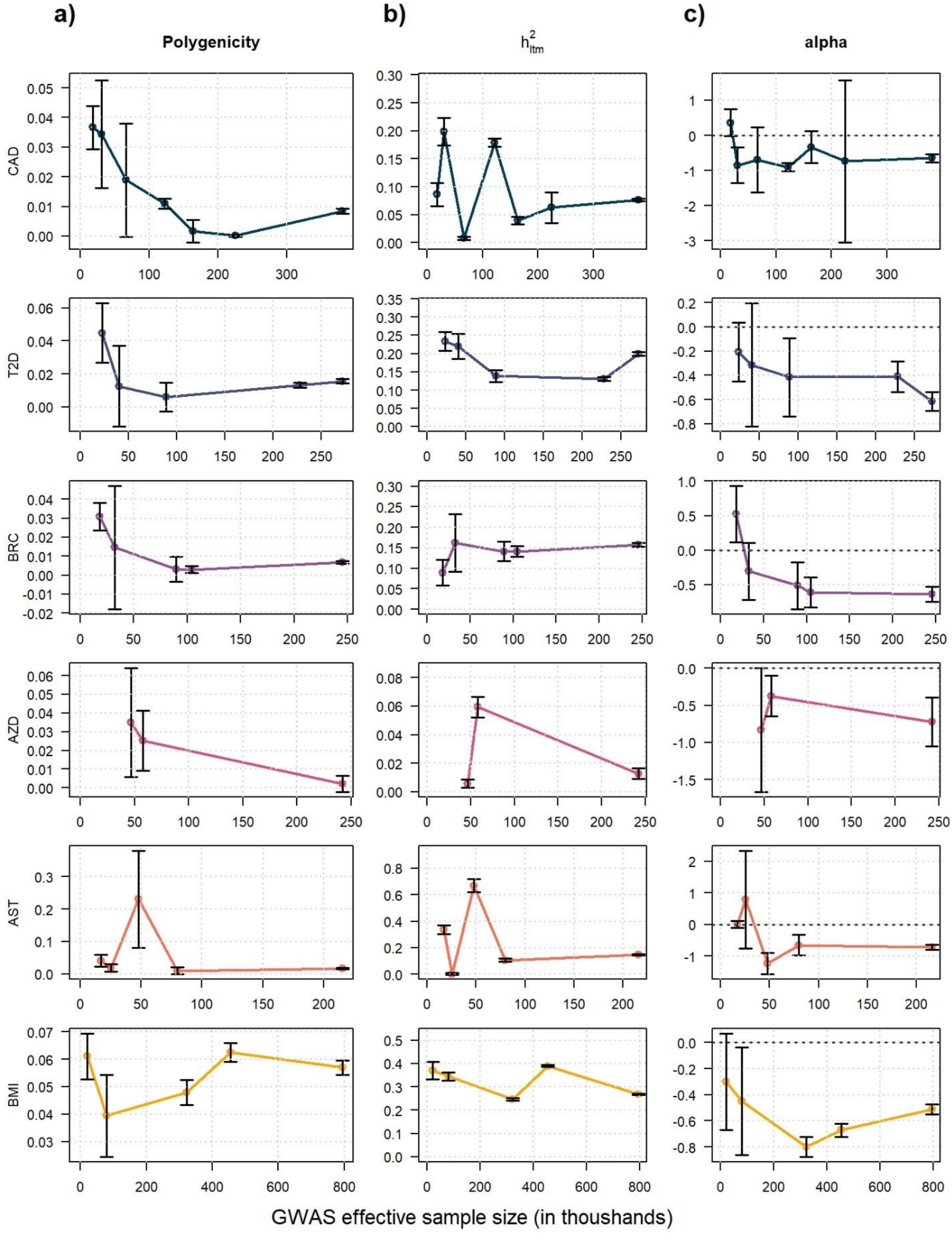

***Figure S8. Comparison between Info_score and true squared-correlation***

Panel a) displays the *info_score* against the squared-correlation between sequenced variants and imputed variants ($r^2$) derived from 125,152 participants of European ancestry from the UK Biobank. Data points are average values derived over bins of equal size of *info_score* values. Panel b) presents the squared-correlation between true genotype and imputed genotype using either a linear additive model (equivalent to the aforementioned $r^2$) or an approximation of the *info_score* derived using genotype probabilities obtained from a multinomial regression. The latter parameters were derived using genotype data from a sample of 20,000 UK Biobank participants of European ancestry.

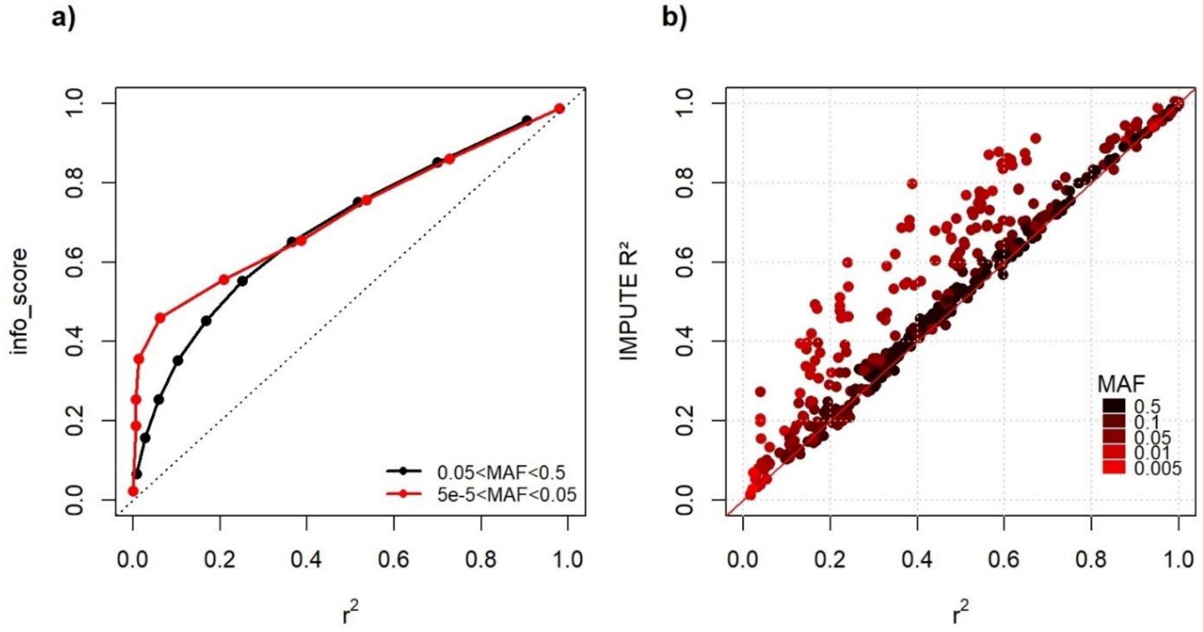

***Figure S9. Variance of untyped variants captured by genotyped variants***

We derived $\rho_j^2$ the variance of each sequenced variant $j$ from the UK Biobank dataset captured by genotyped variants it its vicinity. We first conducted a pilot analysis used genetic data from chromosome 22 to determine the size of the windows around each variant $j$ to be used for an optimal estimation of $\rho_j^2$. For this pilot, we predicted the best guess value from imputed genotypes (referred as the index variant) using genotyped variants as predictors while varying the size of the window in ±[100Kb, 3Mb] around the index. For each index we derived: i) the adjusted r-squared from a multiple regression derived in the entire sample; and ii) the squared-correlation between the predicted index and the true index using a train-test approach, were 66% (N=100K) of the sample was used as a train set and the remaining 33% (N=50K) as a test set. Panel a) shows the average $\rho_j^2$ over all variants of the two metrics for each windows size considered in the pilot study, highlighting an optimal at ±1.5Mb. This optimal window was used in the sequenced data to derive $\rho_.^2$. Panel b) compares the $\rho_.^2$ against $r_.^2$, the squared-correlation between the sequenced variants and the imputed ones in the full chromosome 22 data. The density of the data points is highlighted by a gradient of colours.

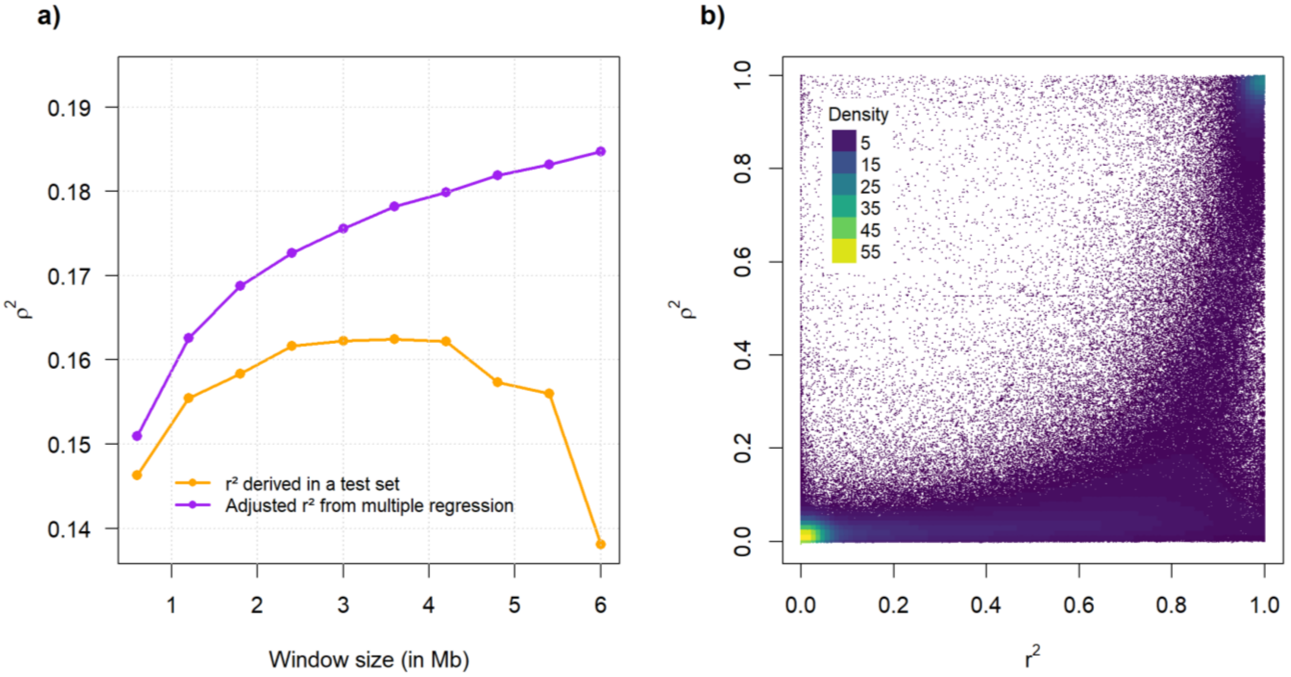

***Figure S10. Attenuated alpha model and proportion of heritability captured***

Panel a) shows the relationship between the minor allele frequency (MAF) and the per-allele effect relative to a base of 1 for a MAF of 0.5 as defined in the $\alpha$ model. Panel b) shows the same relationship for $a = -0.3$ after applying the proposed *ad hoc* attenuated model and using various weights. Panel c) shows the heritability captured by imputed variants and derived based on $r^2$, the squared-correlation between the sequenced variants and the imputed ones, as a function of the value of alpha. Panel d) shows the heritability captured by genotyped variants and derived based on $\rho^2$, the variance of sequenced variants captured by genotyped variants through linkage disequilibrium as a function of the value of alpha. For both estimations we assumed the genetic effect are distributed randomly on the genome based on the $\alpha$ model after applying the attenuation described in panel b.

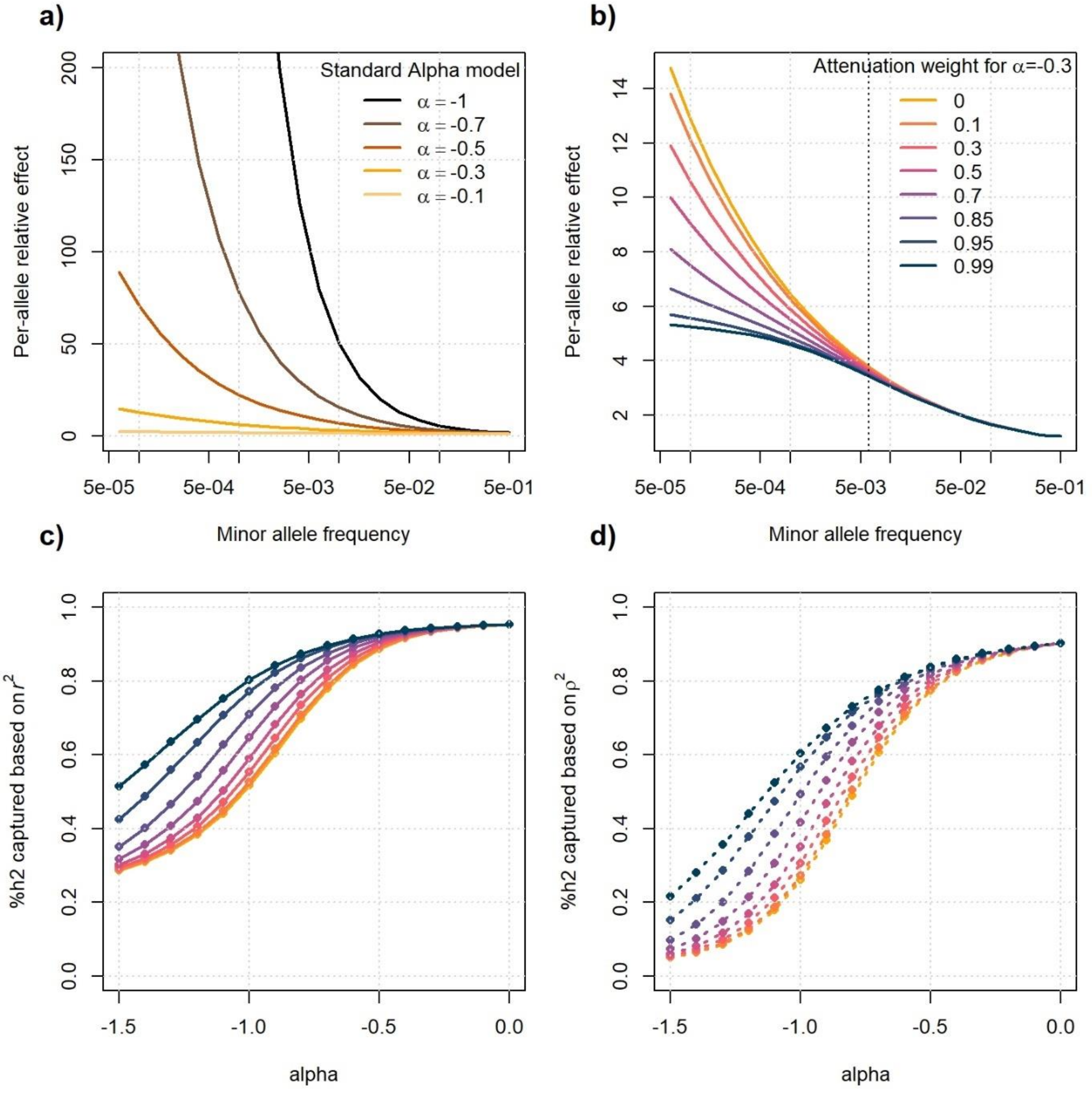

***Figure S11. Estimation of alpha using UK Biobank individual-level data***

Alpha was estimated using individual-level data from UK Biobank participants for the six outcomes: coronary artery disease (CAD), type 2 diabetes (T2D), breast cancer (BRCA), Alzheimer disease (AZ), asthma (AS), and obesity, but also for body mass index for comparison purposes. We considered three models: the standard alpha model (blue plots), the alpha model after applying an attenuation with a weight factor of $w = 0.6$ (red plots), and $w = 0.95$ (green plots), as described in the method section and Figure S9. The panels present the -log(likelihood) as a function of the alpha use to build the GRM (genetic relationship matrix). Panel a) shows the full data, and panel b) a zoom on the y-abscise to highlight the maximum after applying the R *loss()* smoothing function.

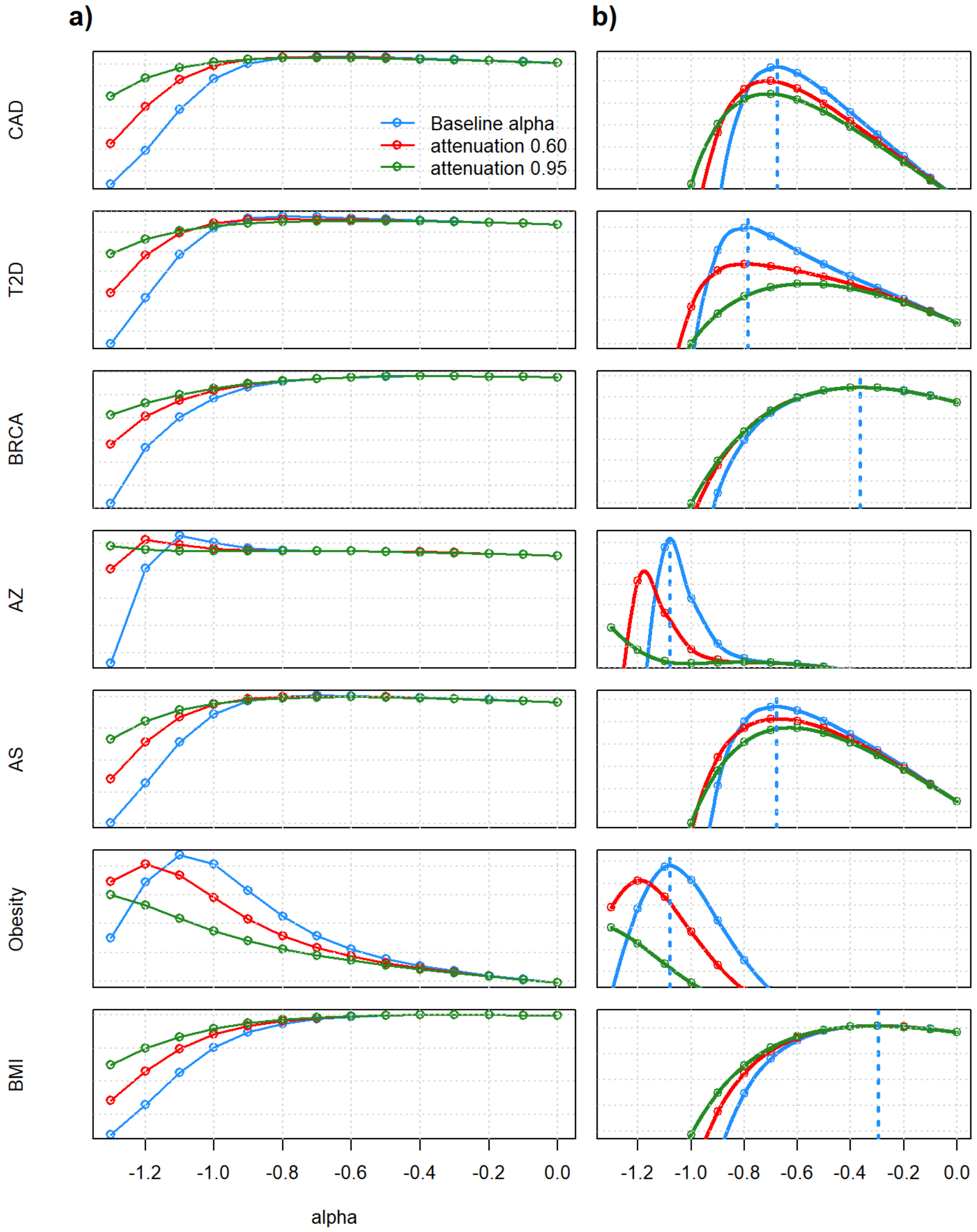

***Figure S12. Validation of the GRM-GCTA pipeline using simulated data***

Estimate of alpha in simulated data using restricted maximum likelihood method as implemented in GCTA. Using 10,000 UK Biobank participants and variants with MAF over 0.1%, we simulated phenotypes using a range of input alpha values. Input alpha values were distributed in the range [-1,0] with a step of 0.1. We then estimated the alpha values using our pipeline, computing the log-likelihood of 11 candidate alpha values in [-1,0] for the simulated phenotypes and selecting the alpha with the maximum likelihood. Results are shown for 10 simulation per input alpha values. The phenotypes were simulated using a heritability of 0.5 and 1% of causal variants randomly drawn from the input data. Both panels correspond to a simulation, the left panel includes all variants available over 0.1% in the simulation, the right panel only includes variants between 0.1% and 1%.

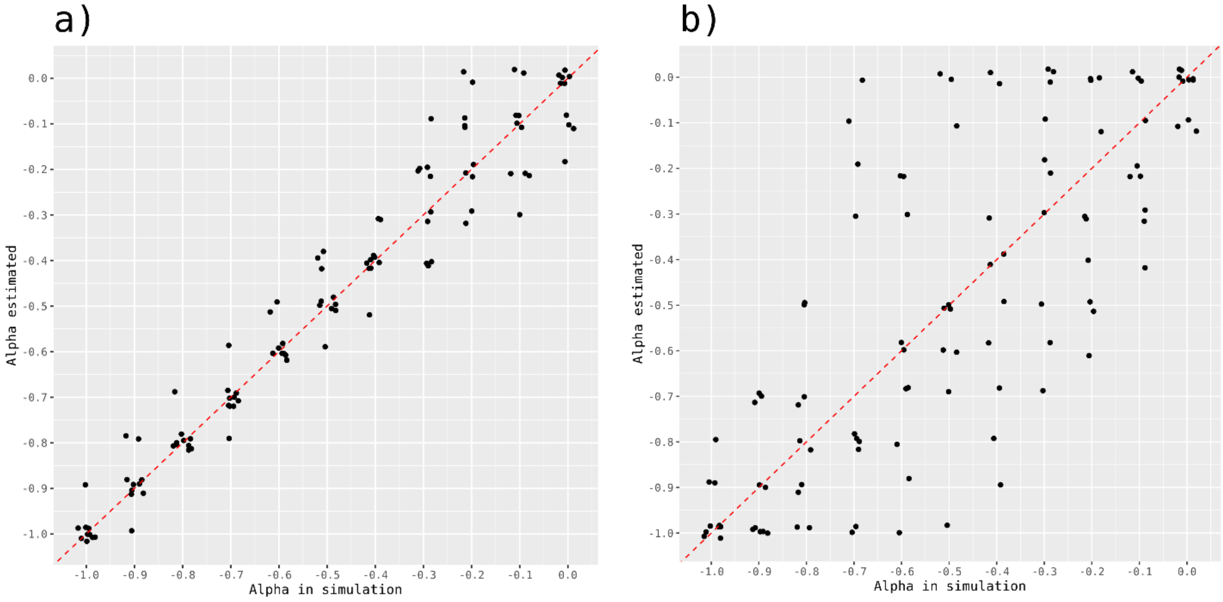

***Figure S13. Correlation between functional annotations***

Correlation between 1,099 functional annotations from nine categories: GENCODE, TFBS (transcription factor binding site), FANTOM5 (functional annotation of the mammalian genome version 5), promoters, enhancers, and dyadic from Roadmap, DHS (DNase I hypersensitive sites) derived from two studies[29], and super enhancer.

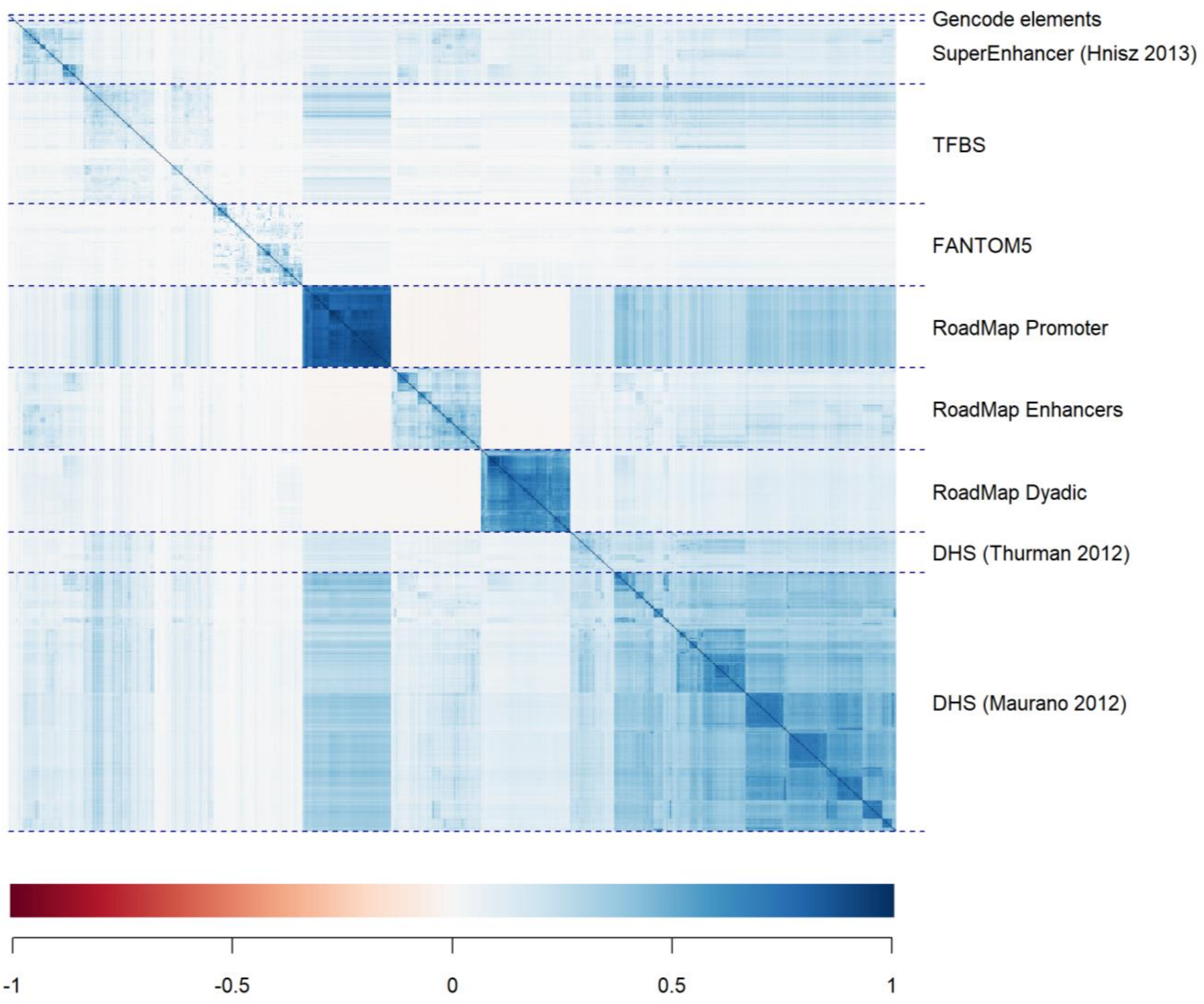

***Figure S14. Distribution of r2 across GENCODE annotations***

Panel a) shows the averaged quality of imputation measured as the squared-correlation ($r^2$) derived between true and imputed genotypes in the UK Biobank for each of the seven GENCODE annotations: intron, gene, exon, CDS (coding DNA sequence), tss (transcription start site), tts (transcription termination site), and UTR (untranslated regions). Panel b) shows the proportion of heritability captured by imputed variants based on $r^2$ and assuming genetic effects are distributed following an alpha model. The relationship was derived assuming the causal variants are randomly distributed (red curve), or assuming the causal variants fall only into each of the seven annotations (green to black gradient).

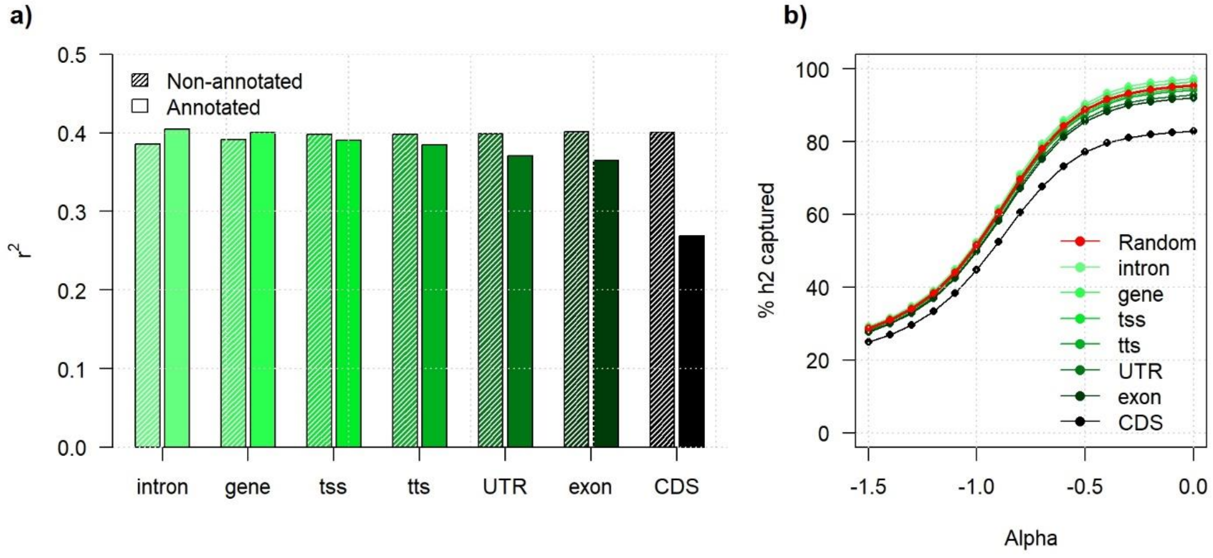

***Figure S15. Top annotations associated with r2***

Relationship between top functional annotations and the quality of imputation measured as the squared-correlation ($r^2$) between true and imputed genotypes. The barplot displays the change in the average $r^2$ between annotated and non-annotated variants for annotation reaching a significance p-value of 1 x$10^{-8}$. The category of each annotation is indicated by a color code: TFBS (transcription factor binding site), FANTOM5 (functional annotation of the mammalian genome version 5), promoters, enhancers, and dyadic from Roadmap, DHS (DNase I hypersensitive sites) derived from two studies, and super enhancer.

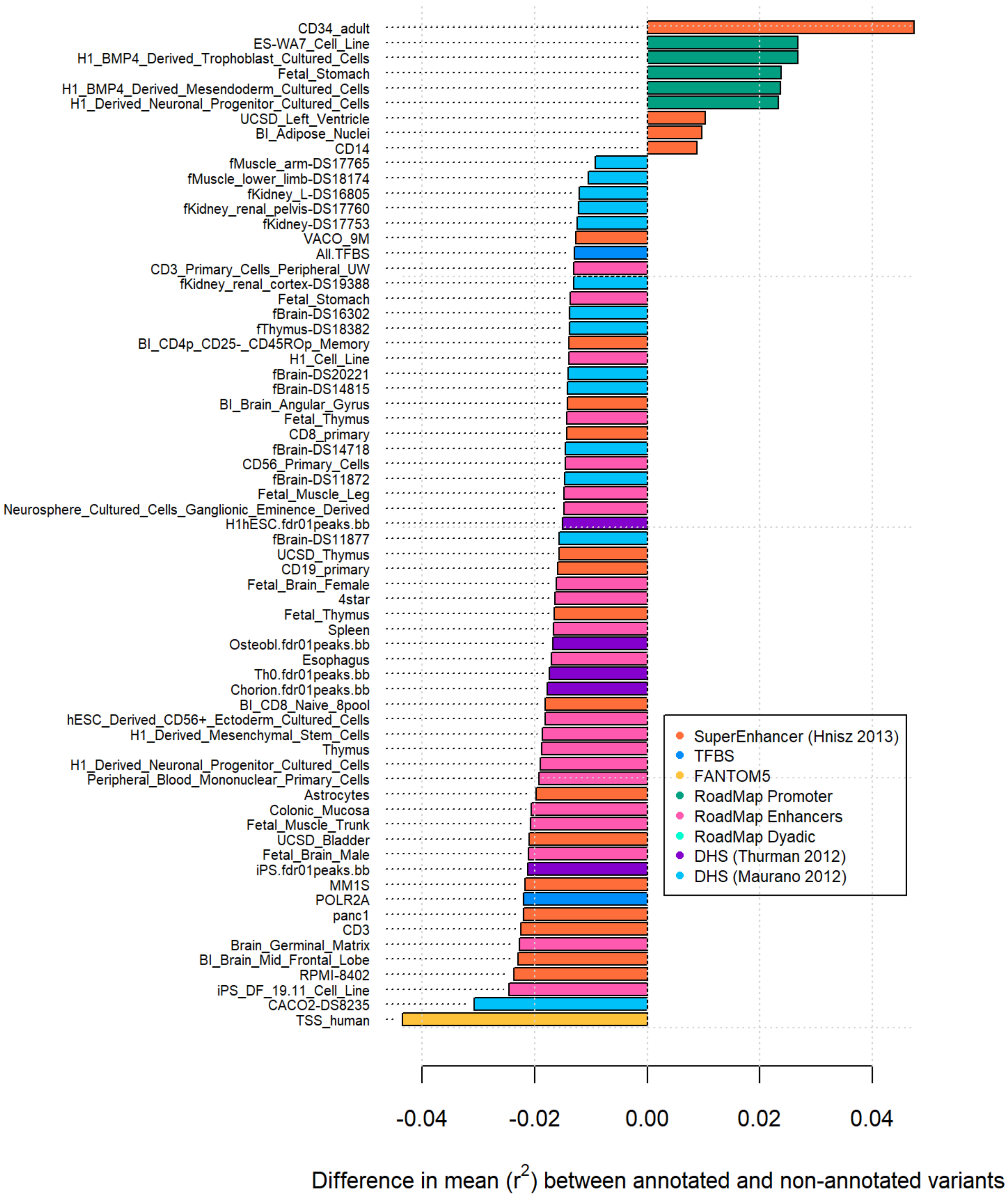
